# Plastic-hydrolytic enzyme classification using explainable deep learning

**DOI:** 10.1101/2025.07.14.664602

**Authors:** Woo-Haeng Lee, Louis Dumontet, KyungMin Jung, Hyun Lee, Gobinda Thapa, Tae-Jin Oh, Mingon Kang

**Affiliations:** Department of Life Science and Biochemical Engineering, SunMoon University, Asan, Republic of Korea 31460; Department of Computer Science at the University of Nevada, Las Vegas, NV, USA 89154; Department of Computer Science and Engineering, SunMoon University, Asan, Republic of Korea 31460; Genome-based BioIT Convergence Institute, Asan, Republic of Korea 31460; Department of Pharmaceutical Engineering and Biotechnology, SunMoon University, Asan, Republic of Korea 31460

**Keywords:** Plastics, Plastic-Degrading Enzymes (PDEs), Plastic hydrolytic enzyme, Classification

## Abstract

The rapid accumulation of plastic waste has emerged as a critical environmental threat, driving the need for scalable and effective biodegradation solutions. Hydrolytic plastic-degrading enzymes (PDEs) offer a promising solution, yet their functional classification remains limited by insufficient annotations and enzymatic diversity. In this study, we present an explainable deep learning framework, PEPIC, to classify nine types of PDEs directly from protein sequences. Using a curated dataset of experimentally validated enzymes and an expanded homologous dataset, we built an explainable deep learning model based on convolutional neural networks (PEPIC) for plastic-degrading enzyme prediction. We benchmarked PEPIC’s performance against state-of-the-art approaches. First, PEPIC demonstrated statistically significant improvements in predictive performance compared to state-of-the-art methods. Second, PEPIC calculates contribution scores for each amino acid in the protein sequence, indicating their influence on the predictions. The model interpretation revealed that regions highlighted by high contribution scores matched conserved catalytic triads and substrate-binding clefts across PET-, PCL-, and PLA-degrading enzymes. Furthermore, structural modeling confirmed the trustworthiness of PEPIC’s predictions. Finally, PEPIC predicted an uncurated enzyme as a PET-degrading enzyme, which was biologically validated to hydrolyze bis(2-hydroxyethyl) terephthalate (BHET). These findings demonstrate that PEPIC provides accurate and trustworthy predictions of PDEs, facilitating the discovery of novel enzymes and supporting the development of sustainable plastic biodegradation technologies.

## 1. Introduction

The environmental crisis caused by the rapid accumulation of plastic waste highlights the need to improve recycling efficiency while maintaining high environmental quality standards.^1–4^ Driven by industrial advancements, global plastic production now exceeds 300 million tons annually. However, due to their inherent durability, these materials do not readily degrade but persist in the environment, causing substantial ecological harm. In particular, polyethylene terephthalate (PET) accounts for nearly 12% of total solid waste and takes up to a century to degrade under natural conditions.^5^ Consequently, extensive research efforts have focused on breaking down synthetic polymers such as PET, polycaprolactone (PCL), and polylactic acid (PLA) into their monomeric components for recycling or upcycling within the framework of circular economy. ^5–7^

Plastic-Degrading Enzymes (PDEs) have emerged as a promising biological solution for the eco-friendly breakdown of synthetic polymers, which attracts significant attention to their structural, catalytic, and ecological diversity.^8–10^ PEDs, often referred to as plastic hydrolases or depolymerases, catalyze the hydrolysis of synthetic polymers into monomers or oligomers, and they facilitate recycling and upcycling processes. Based on their reaction mechanisms, PDEs are broadly categorized into hydrolases and oxidoreductases. Plastic-degrading hydrolases are typically characterized by a conserved α/β-hydrolase fold core and a catalytic triad and directly cleave ester bonds within polymer backbones.^11,12^ On the other hand, xidoreductases, including laccase, peroxidase, and dioxygenase, primarily participate in the oxidative pretreatment of plastic polymers, particularly nonpolar polymers, by modifying polymer surfaces to enhance accessibility for subsequent hydrolytic reactions.^9,13^ Since their activity is often indirect and relies on synergistic interaction with hydrolases, oxidoreductases’ mechanistic roles are difficult to characterize, which limits their usage as direct targets in enzyme engineering and biodegradation studies.

Recent structural and biochemical studies have addressed key features that underlie enzymatic plastic degradation, such as the presence of surface-binding domains (SBDs), hydrophobic patches for polymer interaction, and widened active-site clefts that accommodate polymer chains. For instance, the crystal structure of PETase from *Ideonella sakaiensis* revealed a broadened substrate-binding cleft and unique serine-histidine-aspartate catalytic triad, which facilitate the hydrolysis of PET.^12,14^ These structural characterizations are not only limited to PET-degrading enzymes but also have been observed across other classes of plastic-degrading hydrolases. Similarly, polycaprolactone (PCL) hydrolyzing enzymes often exhibit a lipase-like architecture with a flexible lid domain covering active sites. This lid domain regulates substrates access through conformational switching.^15^ These enzymes typically possess a hydrophobic binding groove that accommodates the aliphatic backbone of PCL, supporting efficient ester bond cleavage. Polylactic acid (PLA)-degrading enzymes also share the α/β-hydrolase fold and possess broad substrate tolerance, typically originating from esterases or lipases.^16^ Structural analysis of PLAases reveals active sites tailored for the aliphatic polyester backbone, often coupled with thermostable scaffolds that facilitate activity under industrial conditions. Such features have been considered to enhance substrate accommodation and catalytic efficiency on polymeric substrates.

Homology-based sequence search has been the primary method to identify candidate enzymes with evolutionary similarity to known plastic hydrolases.^17^ The rapid development of high-throughput sequencing technologies has facilitated large-scale mining of metagenomic and proteomic databases for enzyme discovery.^10,18–20^ However, reliance on sequence similarity alone presents critical limitations on characterizing plastic-degrading enzymes. Functional divergence frequently occurs without significant changes in sequence identity, often hindering the inference of catalytic activity based solely on homology.^10,17,21^ For instance, PETase from *Ideonella sakaiensis* 201-F6 shares only 51% sequence similarity with PET hydrolase from *Thermobifida fusca*, despite having the same function.^22^ Furthermore, phylogenetic approaches, such as Enzyme Commission (EC), do not always align with actual substrate specificity: enzymes from the same EC category may degrade different plastic types, and those degrading the same plastic may fall into distinct EC classes. These inconsistencies pose challenges for both annotation and experimental prioritization of enzyme candidates.

Recent advancements in machine learning have demonstrated promising capabilities for protein sequence analysis. Notable deep learning models for protein sequence analysis include Convolutional Neural Networks (CNNs) and transformers. CNNs effectively capture sequential dependencies of amino acids, which offers strong generalization across homologous proteins and provide interpretable representations.^23–25^ Transformers excel at modeling long-range interactions by capturing structural context, and have achieved state-of-the-art performance when trained with large-scale datasets.^26–28^ Furthermore, explainable deep learning approaches have identified key functional regions in enzyme sequences, which provide trustworthiness in the models’ predictions and potentially guide enzyme engineering.^24,29,30^ This predictive performance and interpretability could accelerate the discovery of novel plastic-degrading enzymes while reducing experimental costs.

The current state-of-the-art framework, Plastic Enzymatic Degradation (PED), identifies plastic-degrading enzymes using XGBoost with transformer-based context-aware enzyme sequence representations.^21^ Despite its successes, PED exhibits several potential limitations. First, the datasets employed for model training and evaluation were relatively small (only 129 sequences belonging to 11 plastic types), which restricts a comprehensive assessment of the model’s robustness across the diverse spectrum of plastic-degrading enzymes. Second, PED considered only the initial 100 amino acids of enzyme sequences, potentially omitting critical functional regions required for accurate enzyme characterization. Lastly, the approach’s reliance on computationally intensive transformers for sequence encoding further restricts its scalability and accessibility.

In this study, we propose PEPIC (Plastic-degrading Enzyme Prediction via Interpretable CNN), which is the first deep learning framework for identifying plastic hydrolytic enzymes from a limited curated dataset (Fig. 1). We first establish a data gathering pipeline to curate high-quality, non-redundant enzyme sequences associated with plastic degradation. Our framework identifies plastic-degrading enzyme sequences and integrates an interpretation module, which highlights the motifs within protein sequences that are most relevant to plastic degradation to provide trustworthy predictions. We benchmarked its predictive performance against transformer-based approaches as well as PED, which is the current state-of-the-art method for plastic-degrading enzyme prediction. PEPIC computes contribution scores of amino acid sequences. We validated that the identified amino acids associated with high contribution scores align with established biological active/binding sites, ensuring the trustworthiness of the predictions. Our findings demonstrated that PEPIC outperforms existing methods for plastic-degrading enzyme prediction and provides interpretability for identifying sequence features associated with plastic degradation. The contributions of this study are: (1) We introduce a data gathering pipeline to assemble a high-quality dataset of plastic-hydrolytic enzymes, utilizing limited biologically annotated data for deep learning-based approaches; (2) we propose an explainable deep learning-based framework to classify plastic-hydrolytic enzymes; (3) we develop an interpretation module to identify motifs relevant to plastic degradation; and (4) we assess that our framework achieves superior predictive performance and produces biologically meaningful interpretations.

**Fig. 1.**
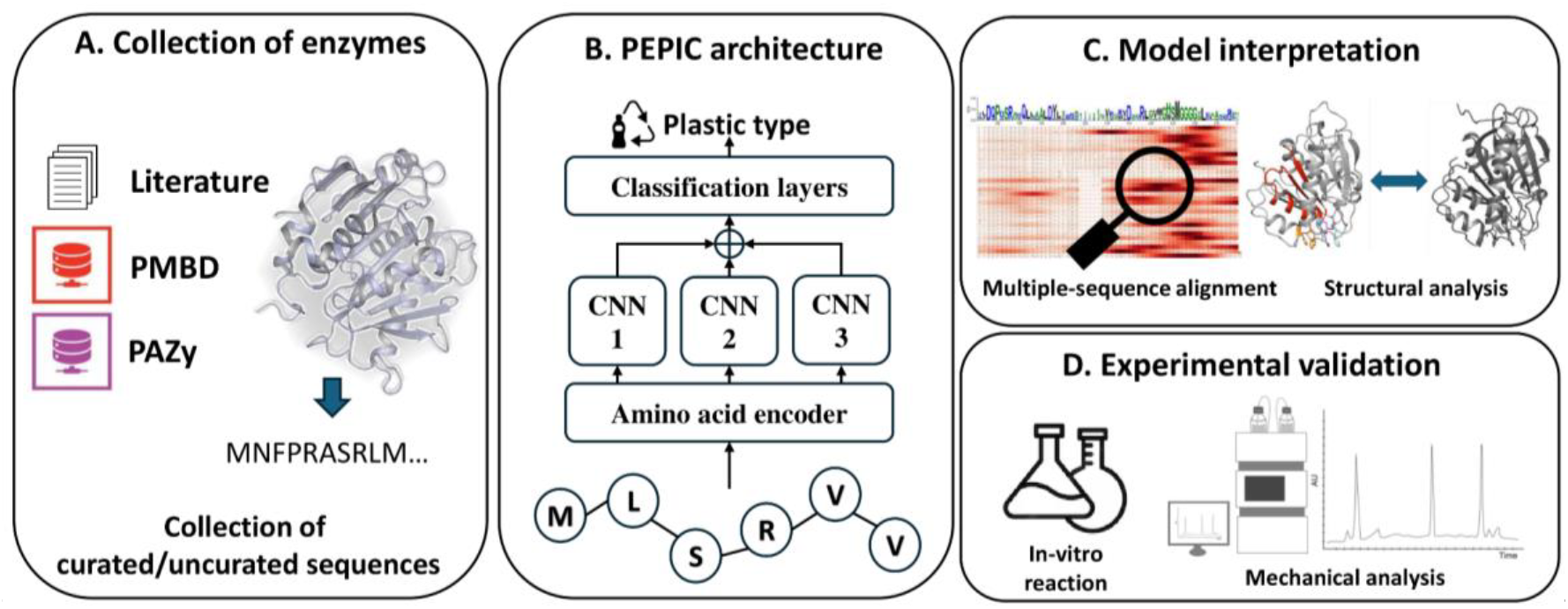
An overview of the study. (A) Collection of protein sequences from PMBD, PAZy, and literature having experimental results. (B) Deep learning architecture of PEPIC for plastic-degrading enzyme prediction. (C) PEPIC provides trustworthy predictions validated by the model’s interpretation by comparing the interpretation with established biological knowledge. (D) Biological experiments for the further validation of an uncurated enzyme candidate. PEPIC identified an uncharacterized sequence, and its plastic-degrading activity was confirmed through biological assays.

## 2. Methods

In this section, we present the architecture of PEPIC, an explainable deep learning framework that predicts plastic-degrading activity from amino acid sequences. We also describe the interpretation strategy to identify sequence regions that are most relevant to the model’s predictions, providing trustworthiness in the predictions.

### 2.1. Architecture of the proposed model

PEPIC analyzes amino acid sequences and predicts enzymes’ plastic-degrading activity. PEPIC employs a convolutional neural network backbone to capture local sequence dependencies, ensure robust generalization across enzyme families, and facilitate interpretability. PEPIC is designed based on the DeepEC for plastic hydrolytic enzyme prediction.^23^ PEPIC consists of (1) a protein sequence encoder, (2) three parallel convolutional layers, (3) a pooling layer, and (4) three sequential classification layers. First, the input protein sequences are encoded using an embedding approach (e.g., one-hot encoding, BLOSUM62 matrix, and ProtVec embeddings). Second, three convolutional layers are applied in parallel to the encoded sequence, with kernel sizes of (4×*d*), (8×*d*), and (16×*d*), where *d* represents the dimension of the selected embedding. Each convolutional layer consists of 128 filters with the ReLU activation function. Third, a 1-max pooling operation is applied to each convolutional output to retain the most prominent features. The pooled outputs are then concatenated into a single 384-dimensional feature vector. Finally, this vector is passed through three fully connected layers of sizes 512, 512, and 11, respectively. The first two fully connected layers are activated by the ReLU function, while the final classification layer uses the sigmoid activation to accommodate the multi-label nature of the task. Model parameters are optimized by minimizing the binary cross-entropy loss between the predicted outputs and the ground truth labels.

### 2.2. Interpretation strategy

We compute contribution scores for individual amino acids to provide domain-specific evidence, ensuring trustworthiness in the predictions. The contribution scores identify protein regions, such as conserved motifs or functional domains by propagating contribution scores from the model’s activation maps. We adapted the explainable deep learning approach developed for CNNs.^24^ Activation maps are generated by convolutional filters of different sizes (*l* = 4, 8, or 16), each capturing patterns of varying lengths. Each activation map produces a sequence of activation values *Φ*_*i*_^*k*^, where each value corresponds to a window of *l* consecutive amino acids. Activation maps are ranked based on their contribution to the final prediction (*α*_*m*_), which is computed as the partial derivative of the prediction score (y_*c*_) with respect to the activation maps’ max-pooled value (*β*_*m*_), such as:

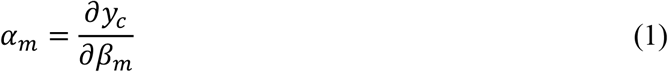

The top *K* activation maps (e.g., *K* = 50) with the highest scores are selected, and their activation values are summed to compute the contribution score (*ζ*_*l*_) for each window:

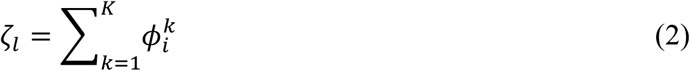

Each activation covers multiple amino acids, its contribution is distributed across individual residues by averaging across all overlapping windows containing amino acid *i*, yielding the contribution score (τ_*i*_^*l*^):

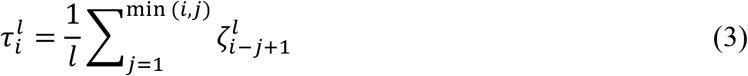

Finally, contribution scores from different filter sizes are aggregated to obtain the final amino acid relevance score:

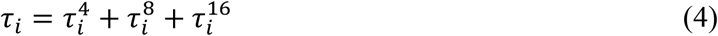

By propagating contribution scores from high-level model activations back to the sequence level, this approach provides an interpretable mapping of functional residues that contribute to a prediction.

## 3. Results

This section first outlines our data gathering pipeline that constructs datasets for model development and evaluation of plastic-degrading enzymes. Second, we assess the predictive performance of PEPIC by comparing it with state-of-the-art benchmark models. Third, we evaluate the trustworthiness of the predictions by aligning PEPIC’s interpretation with established biological knowledge. Finally, we assess the robustness of PEPIC through biological assays on an uncharacterized enzyme candidate predicted by the model.

### 3.1. Dataset collection for model development

We constructed plastic-degrading enzyme datasets for deep learning model development and evaluation. In this study, we focused on hydrolytic enzymes, a well-characterized subclass of plastic-degrading enzymes. We excluded oxidoreductases, due to limited data availability and unclear mechanistic relevance in the context of plastic hydrolysis. First, we collected biologically curated hydrolytic enzyme sequences through a literature review. Then, we generated a model development dataset using sequence similarity between UniProt and the biologically curated dataset.

#### 3.1.1. Curated datasets from literature and PlasticDB

We gathered biologically curated datasets from both peer-reviewed literature and the PlasticDB.^31^ We surveyed literature from PubMed published before April 2022 with the following search query: [Plastic* AND (*degrad* OR depolymer*) AND (bacter* OR fung* OR archaea*)]. We manually examined the queried papers that reported experimentally validated plastic-degrading enzymes. Then, we confirmed whether the enzymes had been validated using both biochemical and analytical methods. We excluded enzyme candidates validated by only weight loss measurements, since polymer mass loss can be caused by abiotic processes, such as photodegradation, oxidation, or surface erosion.^8^ We also cross-referenced the manually curated dataset with PlasticDB (https://plasticdb.org, accessed June 2023). From PlasticDB, we selected the sequences annotated as hydrolytic enzymes based on both UniProt functional annotations and InterproScan-predicted domains, including alpha/beta-hydrolase, esterase, and cutinase, which ensures domain-level evidence for potential plastic-degrading function. In the resulting dataset, the plastic-degrading activities were categorized into the nine plastic types: PHA (polyhydroxyalkanoate), PHB(Polyhydroxybutyrate), PBAT(poly(butylene adipate-co-terephthalate)), PBS(poly(butylene succinate)), PBSA(poly(butylene succinate-co-adipate)), PLA (polylactic acid), PCL (Polycaprolactone), PET (poly(ethylene terephthalate)) and PU (polyurethane). Finally, we obtained a biologically curated dataset comprising 181 non-redundant plastic-degrading enzyme sequences. This dataset consists of 50 PHA-degrading, 46 PHB-degrading, 16 PBAT-degrading, 11 PBS-degrading, 16 PBSA-degrading, 26 PLA-degrading, 40 PCL-degrading, 75 PET-degrading, and 14 PU-degrading enzymes and was used to evaluate the predictive performance of the models.

#### 3.1.2. Model development dataset

We derived a large training dataset from the biologically curated dataset using sequence similarity for model development. Specifically, we identified potential plastic-degrading enzymes in UniProt by performing BLASTp^32^, using the enzyme sequences in the biologically curated dataset as queries (bit-score ≥ 100 and sequence identity ≥ 85%). Additionally, we excluded sequences with large length discrepancies compared to the query enzymes to maintain functional and structural comparability as well as minimizing contamination from non-target domains. To reduce data redundancy, we applied CD-HIT clustering with a 90% identity threshold^33^, which resulted in a final positive dataset of approximately 5,927 enzyme sequences. The model development dataset consisted of 253 PBAT-degrading, 75 PBS-degrading, 469 PBSA-degrading, 1,367 PCL-degrading, 1,473 PET-degrading, 2,657 PHA-degrading, 1,804 PHB-degrading, 1,195 PLA-degrading, and 313 PU-degrading enzymes, and the dataset was used for cross-validation.

We constructed a negative dataset of non-plastic-degrading enzymes belonging to the broader α/β-hydrolase superfamily, which includes enzymes that are structurally similar but not associated with plastic degradation. We obtained the negative dataset from the Lipase Engineering Database (LED)^34^, ensuring coverage of carboxylesterases, lipases, and other related enzymes. To prevent overlapping with the positive dataset, we removed all sequences that exhibited above 80% identity similarity with either the biologically curated or model development datasets. Finally, the negative dataset comprised 30,350 enzyme sequences. The negative dataset was used for computing false discovery rate (FDR).

### 3.2. Predictive performance comparison using cross-validation and biologically curated data

We evaluated the performance of PEPIC to predict plastic hydrolytic enzymes by comparing it against transformer-based models and PED.^21^ We considered an encoder-only transformer^35^, which is the typical transformer architecture for protein sequence analysis. The details of the transformer architecture are provided in Supplementary Note 1. For this benchmark experiment, we split the model development dataset into training (80%), validation (10%), and test (10%), with stratified sampling to preserve the class ratios. We optimized the models with the training set, and the hyper-parameters were fine-tuned with the validation set. The protein sequence encoder was optimized using the validation set among thirteen encoding schemes^36^, the detailed results are presented in the Supplementary Note 2. The optimal hyper-parameter values of PEPIC were a learning rate of 1e-4, a dropout of 0.1, and protein sequences were encoded with Miyazawa energies. PEPIC was trained using the ADAM optimizer. The optimal hyper-parameter values of the transformer model used for comparison were a model dimension of 256, 2 attention heads, 4 encoder layers, a dropout rate of 0.1, and a learning rate of 1e-4. The transformer model was also trained using the ADAM optimizer. For PED, we used the hyper-parameter values reported in the original study.^21^ Note that its predictive performance has been shown to be relatively non-sensitive to hyper-parameter variations due to the strong inductive bias introduced by the large-scale pre-trained transformer it employs. We assessed the predictive performance of all models on the test dataset by computing the micro-averaged F1-score, precision, and recall. We selected the optimal thresholds for the discriminative function that maximize micro-averaged F1-score on the validation set. We also reported the false discovery rates (FDR) computed from amino-acid sequences in the negative dataset. We used the same training, validation and test datasets for all benchmark models, and the experiment was repeated twenty times for reproducibility. PEPIC significantly outperformed transformer-based models and PED on the cross-validation dataset (Fig.2A). PEPIC achieved an F1-score of 0.989±0.098, precision of 0.999±0.001, and recall of 0.979±0.019, while the transformer model obtained 0.965±0.030, 0.989±0.011, and 0.954±0.044, respectively. PED achieved an F1-score of 0.679±0.044, precision of 0.677±0.029, and recall of 0.726±0.036. This corresponds to an improvement of approximately 2.5% in F1-scores over the transformer model and 44% over PED. The superiority of PEPIC over the benchmark models was statistically validated using a Wilcoxon signed-rank test, yielding a p-value of less than 0.01.

We applied the twenty models optimized on the cross-validation dataset to the biologically curated dataset to further evaluate their generalization performance (Fig. 2B). On this biologically curated dataset, PEPIC achieved an F1-score of 0.757±0.018, precision of 0.875±0.031, and recall of 0.670±0.040. The transformer model obtained an F1-score of 0.661±0.056, precision of 0.811±0.035, and recall of 0.561±0.070, while PED achieved 0.624±0.053, 0.623±0.012, and 0.701±0.025, respectively. These results correspond to an improvement of approximately 15% over the transformer model and 21% over PED in F1-scores. On the negative dataset, the FDR values were 0.029 ± 0.012 for PEPIC, 0.044±0.013 for the transformer, and 0.057±0.001 for PED.

**Fig. 2.**
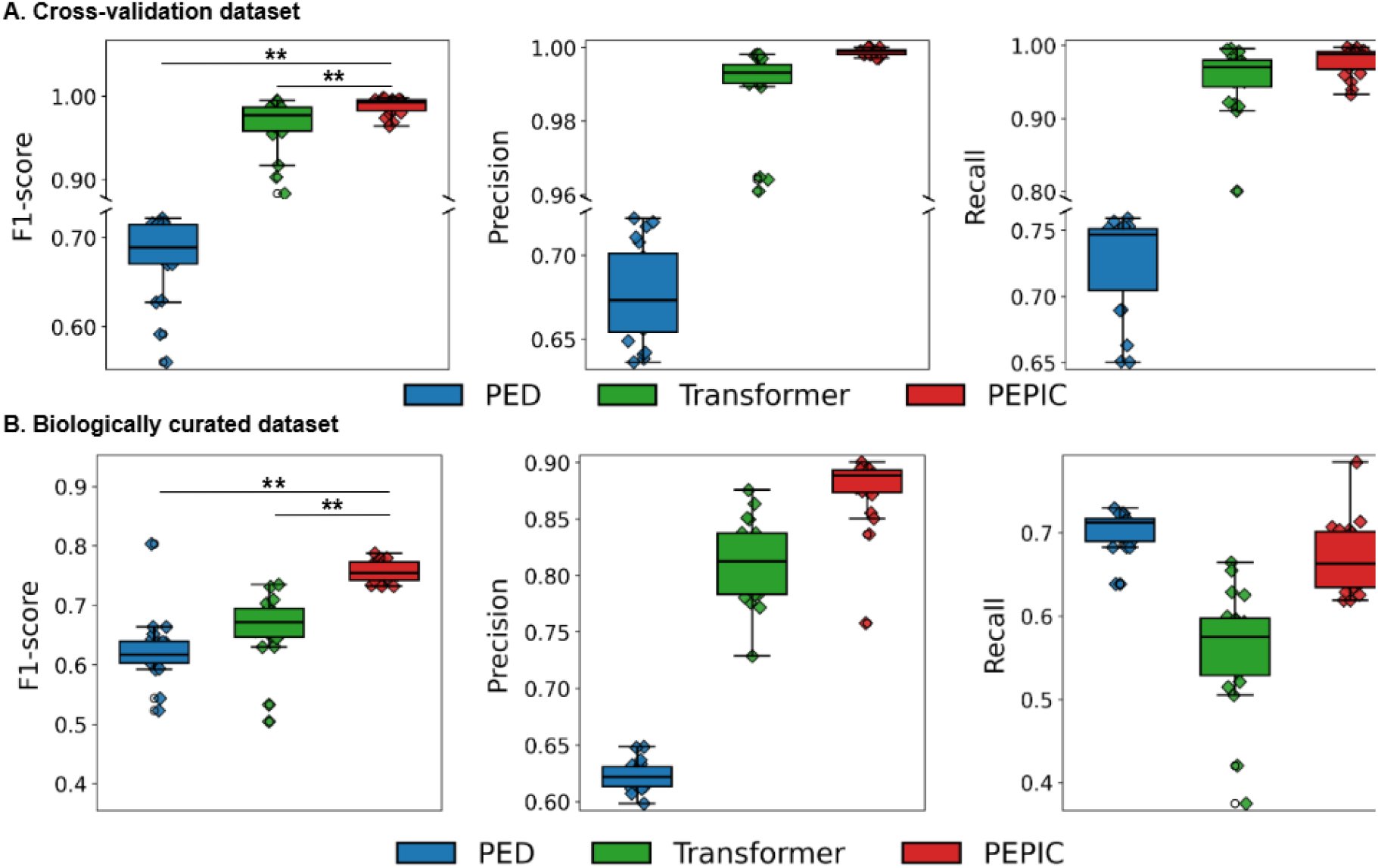
Predictive performance of PED, transformer, and CNN (A) on the cross-validation dataset and (B) biologically curated dataset. ** indicates statistically significant improvements based on the Wilcoxon signed-rank test (p < 0.01).

### 3.3. Biological validation of PEPIC’s interpretation

We evaluated the biological relevance of PEPIC’s contribution scores to support its trustworthy predictions by comparing high-contribution sequence regions with established functional motifs in plastic-degrading enzymes. For a given protein sequence, we computed contribution scores for each amino acid to identify the most influential regions to the model’s predictions. For the assessment, we randomly selected PET-, PCL-, and PLA-hydrolyzing enzymes, and computed their contribution scores (Fig. 3). Then, we compared the most contributing residues with conserved motifs from well-characterized references. We further performed homology modeling to map these residues onto three-dimensional structures, confirming their localization within catalytically important regions. All the sequences that we used are in Supplementary Table 1.

**Fig. 3.**
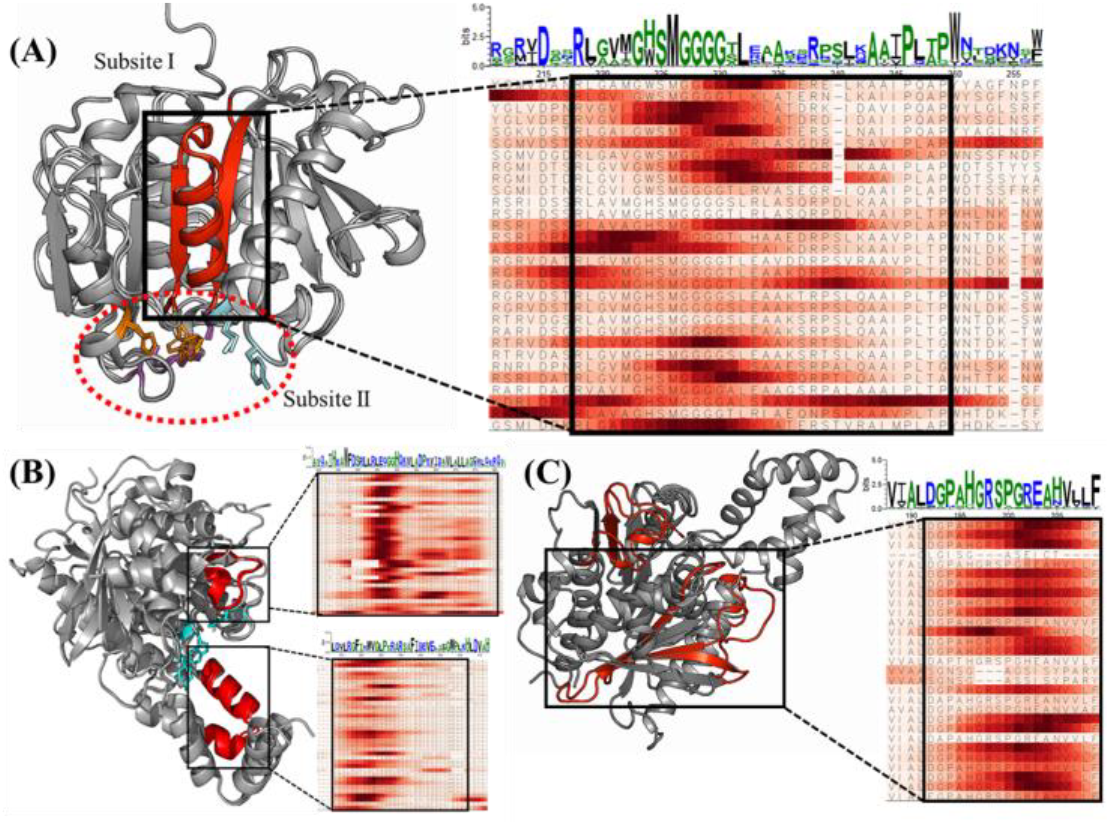
Interpretation results of model-derived importance and structural conservation of a key motif in hydrolytic plastic-degrading enzymes. (A) Distribution of importance score and structural analysis of PET-degrading enzyme, (B) Distribution of importance score and structural analysis of PCL-degrading enzyme, (C) Distribution of importance score and structural analysis of PLA-degrading enzyme.

For PET-hydrolases, PEPIC showed high contribution scores in subsite I that includes the nucleophile elbow (Gly-X-Ser-X-Gly motif) (Fig. 3A). The serine residue of the nucleophile elbow plays a critical role in cleaving ester bonds and is a conserved residue among PET-hydrolases. Subsite I, composed of an α-helix and β-sheets, is a part of the α/β-hydrolase fold, as conserved structural features. Specifically, Met226 and Trp/His224 are involved in interactions with the PET polymer, and PEPIC identified them with high contribution scores. Trp260 contributes to substrate binding affinity through π-π interactions with the aromatic rings of PET.^12^

For PCL-hydrolases, PEPIC produced high contribution scores around histidine among the catalytic triads. Specifically, high contribution scores were assigned to the histidine-containing loop (upper box in black) and the helix parts (bottom box in black) in Fig. 3B. The histidine-containing loop has high flexibility, which allows flexible adaptation of the substrate binding cleft of the enzyme, thereby increasing substrate specificity.^37^ This helix part is a variable structure covering the active site observed in lipases^15^ and is directly related to substrate specificity, active site accessibility, and reaction mechanism.

For PLA-hydrolases, PEPIC identified several loops and sheets that are distant from the catalytic site (Fig. 3C), as well as histidine-containing loops like PCL-hydrolases. The loops and sheets can be interpreted as structural adaptations to accommodate flexible, aliphatic substrates. Unlike PET-hydrolyzing enzymes, most PCL or PLA-hydrolyzing enzymes form a pocket-shaped structure to bind aliphatic substrates.^16,38,39^ In particular, the β4-α3 loop is significantly extended compared to PET hydrolases’ ones, contributing to the formation of a substrate-binding pocket. Similarly, the β3-β4 loop and histidine-containing loop are known as contributing to shaping the substrate-binding pocket and determining substrate specificity.

### 3.4. External validation with an uncurated enzyme

We evaluated the robustness and generalization capacity of PEPIC by applying it to a plastic-degrading candidate enzyme that has no prior experimental annotation. To select the best candidate enzymes in this experiment, we performed a local BLAST search on microbial genomes isolated from extreme environments (e.g., psychrophilic or photosensitizer-associated strains), querying PETase from *Ideonella sakaiensis* (UniProt ID: A0A0K8P6T7). Among them, we found a candidate gene exhibiting low sequence identity (i.e., similarity < 40%) and unclear functional annotation but having conserved domain in multiple-sequence alignment with known PET-hydrolases. We investigated both its enzymatic activity and the interpretability of the model’s prediction to determine the model’s reliability in identifying novel plastic-degrading enzymes beyond curated datasets. PEPIC predicted the enzymes’ PET-hydrolyzing activity with a probability of 59.3%. To verify the prediction biologically, we heterologously expressed and purified the enzyme (W2061_PET9), followed by *in-vitro* enzymatic assays using bis (2-hydroxyethyl) terephthalate (BHET) as a substrate, which is one of the intermediate compounds in PET degradation. In the biological experiment, this enzyme successfully cleaved the ester bond in BHET during *in-vitro* assays, producing Mono(2-hydroxyethyl) terephthalate (MHET) as the product (Fig. 4A). HPLC and LC-MS analysis of the reaction mixture showed a substrate peak at around 11 minutes and a product peak corresponding to MHET at 10.5 minutes, with molecular weights of m/z 287.0537 and m/z 227.9563, respectively, which confirms the function of PETase (Fig. 4B). Furthermore, the interpretation for this candidate enzyme was consistent with those of known PET-hydrolases. As shown in Fig. 5, high contribution scores were observed in subsite I, including the nucleophile elbow. Although the overall score was slightly lower than that of experimentally validated enzymes, the high contribution in the nucleophile elbow suggests that PEPIC effectively identified functionally relevant features.

**Fig. 4.**
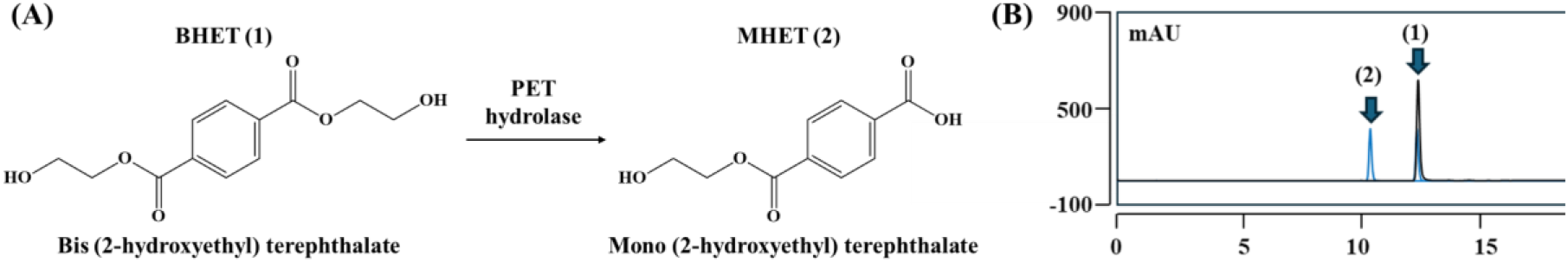
Results of the enzymatic PET-hydrolysis: (A) Scheme of enzymatic PET-hydrolysis reaction, (B) HPLC separation of the mixtures of *in-vitro* reaction for W2061_PET9 with Bis(2-hydroxyethyl) terephthalate. (1) Mono (2-hydroxyethyl) terephthalate, (2) Bis(2-hydroxyethyl) terephthalate.

**Fig. 5.**
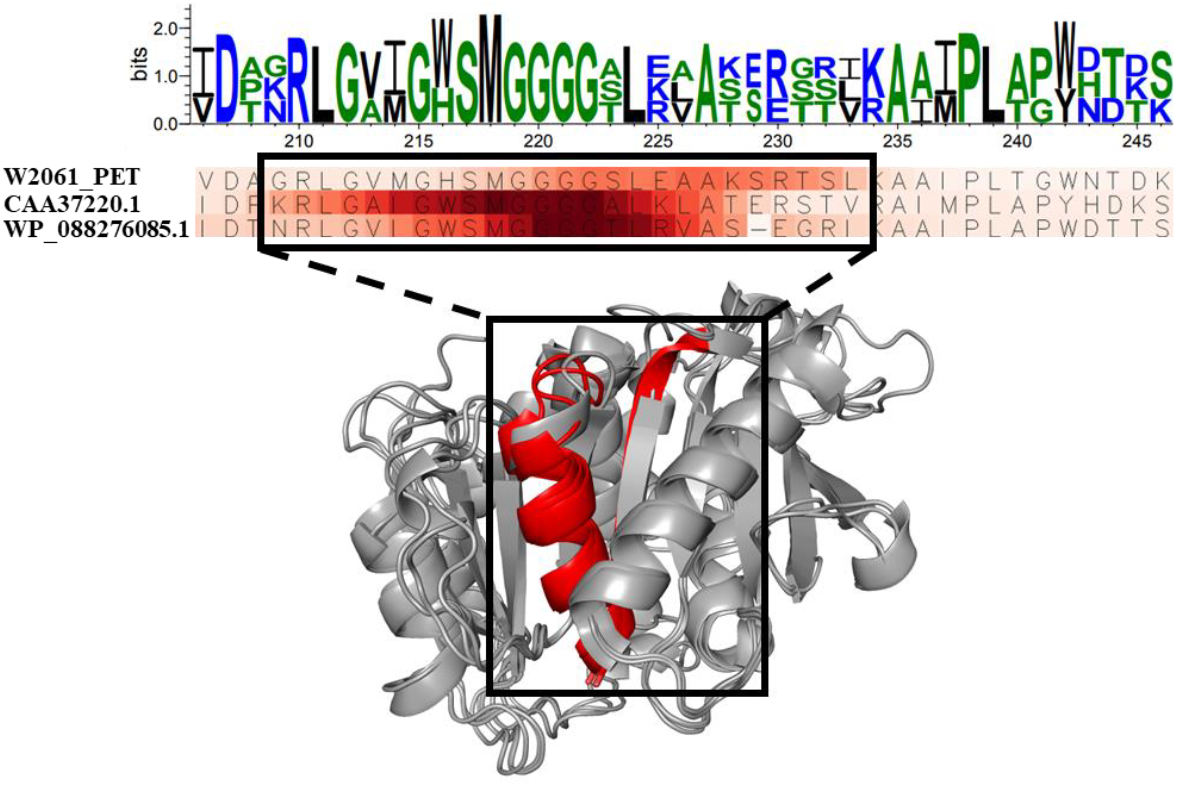
Interpretation results of model-derived importance and structural conservation of a key motif in hydrolytic plastic-degrading enzymes. (Top) Sequence logo generated from a multiple sequence alignment of hydrolytic plastic-degrading enzymes, highlighting conserved residues within the 210-245 amino acid region. (Middle) Heatmap visualization of residue-level importance scores derived from the trained deep learning classification model. Shades of red indicate higher attention weights assigned by the model, with the GXSXG motif showing consistently elevated importance across sequences. This region overlaps with highly conserved residues identified in the sequence alignment. (Bottom) Structural mapping of the high-importance region onto the reference enzyme structure. The red-highlighted segment corresponds to the GXSXG motif and its adjacent residues, positioned near the substrate-binding pocket. The alignment of model-derived attention with structural and evolutionary conservation suggests a potential role in substrate recognition and catalytic specificity.

## 4. Discussion and conclusion

In this study, we have developed a CNN-based explainable deep learning framework (PEPIC) for accurately classifying hydrolytic plastic-degrading enzymes (PDEs). The model outperformed existing machine learning and deep learning approaches, demonstrating high power of predictive performance on both cross-validation and curated biological datasets. Importantly, PEPIC provides robust model interpretability for trustworthy predictions. Our interpretation revealed that PEPIC effectively recognizes biologically meaningful regions associated with substrate binding and catalytic activity. These results support that the PEPIC can capture evolutionarily conserved and structurally significant patterns essential for enzymatic plastic hydrolysis. The biological validation and assessment of the proposed approach address its potential to classify novel PDEs based on active-site architectures in a substrate-specific manner.

However, this study may have potential limitations and discussion. First, the high predictive performance on the cross-validation may be produced due to the high sequence similarity on the datasets. Nevertheless, PEPIC maintained superior performance on the biologically curated dataset that presents more sequence diversity and less redundancy, which demonstrates its robustness. In contrast, transformers exhibited intermediate performance across both datasets. Transformers typically require larger datasets to fully leverage their capacity for modeling long-range dependencies, and the performance is expected to improve with access to broader and more diverse training data. Overall, PEPIC is highly effective for plastic hydrolytic enzyme prediction in settings with limited curated data.

Second, incorporation of oxidoreductase data and diverse plastic types are required to improve PEPIC’s utility for comprehensive plastic biodegradation research and sustainable biotechnology applications. We focused only on hydrolytic plastic-degrading enzymes, mainly due to the limited availability of oxidoreductase data. However, enzymes, such as laccases and peroxidases, also play important biological roles in plastic degradation. Additionally, our training set did not include enzymes that degrade aliphatic plastics without ester bonds, such as polystyrene (PS) or polyvinyl chloride (PVC). Expanding the dataset to include these enzyme types and substrates is essential for broader applicability.

This work provides a reliable computational tool to accelerate the discovery of plastic-degrading enzymes, contributing to sustainable plastic waste management. Further investigation, including expanding enzyme diversity and incorporating additional polymer targets, would be expected as future work.

## Acknowledgements

This work was supported by the National Science Foundation Major Research Instrumentation (NSF MRI) (Grant#:2117941), the Institute of Information & Communications Technology Planning & Evaluation (IITP) funded by the Korean government (MSIT) (No. 2021-0-01581), and the Bio & Medical Technology Development Program (RS-2024-00441423) funded by the National Research Foundation of Korea (NRF).

## Author Contributions

W-H.L and L.D. conceived the original idea, designed and conducted the experiments, and wrote the manuscript. K.M.J and H.L contributed computational experiments. G.T. conducted biological experiments for the assessment. T-J.O. conceived the original idea, provided critical revisions, secured funding for the project, and oversaw the overall direction of the work. M.K. supervised the project, contributed to manuscript writing and revisions, provided funding, and oversaw the overall direction of the work. All authors have reviewed and approved the final manuscript.

## Data Availability

Protein sequences used for biological assessment are available in Supplementary Table 1. All other relevant data are available from the authors upon request.

## Code Availability

The open-source code is available at: https://github.com/datax-lab/Plastic

## Ethics Declarations

The authors declare no competing interests.

