## Supplementary for "Plastic-hydrolytic enzyme classification using explainable deep learning"

**Supplementary Note 1. Transformer architecture.**

We considered an encoder-only transformer for the benchmark. It is composed of an embedding module followed by four encoders. The embedding module relies on two layers: the amino acid matrix embedding and the positional encoding. The amino acid matrix embedding is a $23 \times d_{model}$ ($d_{model} = 256$) matrix learned during training such that the amino acid (assigned index $a$) has its embedding in the $a$-th row of this matrix. Note that we need an embedding for the 20 different amino acids: the undetermined amino acid often referred to X, the classification token, as well as the padding token which account for the 23 rows of our matrix. The positional encoding module takes the position $p$ of the amino acid as input and encodes it as:

$$PE(p, 2i) = sin(\frac{p}{10000^{2i/d_{model}}}), \tag{1}$$

$$PE(p, 2i + 1) = cos(\frac{p}{10000^{2i/d_{model}}}), \tag{2}$$

where 2i and 2i+1 represent even and odd indices, respectively. The outputs of the two layers are then added to form the output of our embedding module, which is fed to the first encoder.

Each encoder comprises a multi-headed attention module followed by two fully connected layers activated by the ReLU function, a dropout layer, and a layer normalization. The multi-headed attention layer consists of projecting the input into 3 vectors $Q$ (query), $K$ (key), and $V$ (value) of size $d_k = 256$. The output of the attention module is computed as:

$$Attention\ Scores = Softmax(\frac{Q \cdot K^T}{\sqrt{d_k}}), \tag{3}$$

$$Attention\ output = Attention\ Scores \cdot K. \tag{4}$$

Then, the output is fed to a dropout layer with a dropout rate of $0.1$. The multi-headed attention module incorporates a skip connection, which adds the input of this module to its output. The given result is normalized and fed to two successive position-wise feed-forward layers of size $4 * d_k = 1024$ in which the first one uses ReLU as an activation function, and both layers apply a dropout of 0.1 to their output. These two layers are skipped through a skip connection that adds the module's input to its output.

The final output of the encoder is passed through a dropout layer with a dropout rate of 0.1. After going through the embedding module and the four encoders, the final prediction is made by a linear layer, which takes as input the representation of the classification token by the last encoder of the transformer and uses the sigmoid function as activation to produce a score between 0 and 1 for each of the 11 classes.

**Supplementary Note 2. Predictive performance comparison of CNNs across various encoders.**

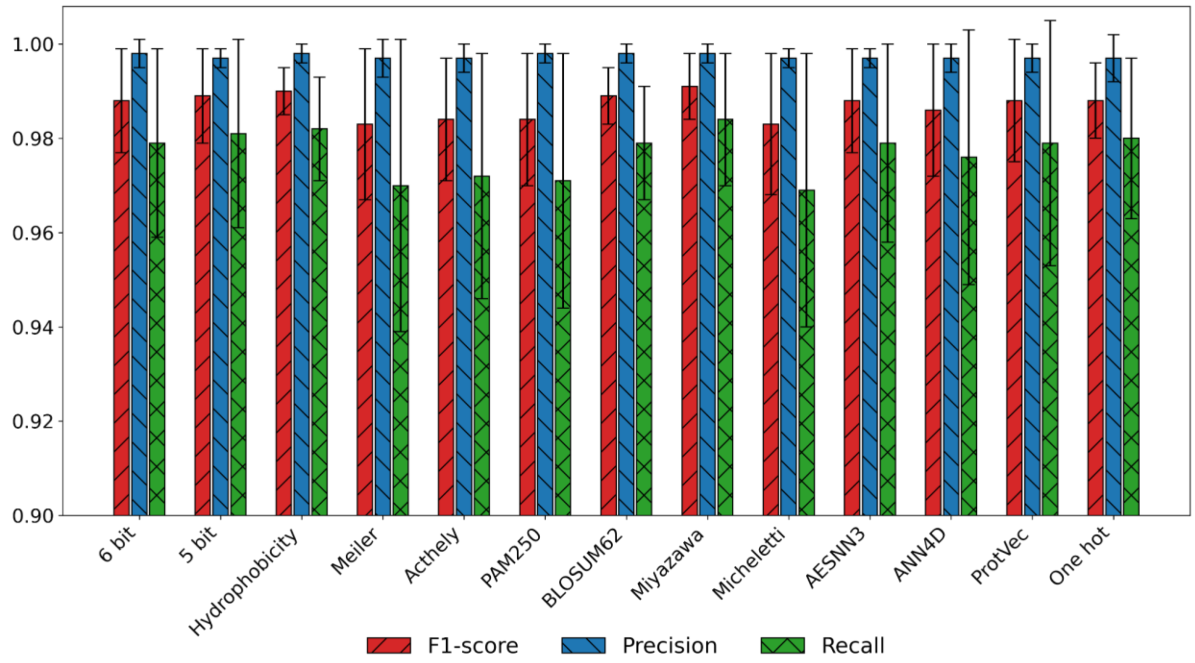


Fig. S1. Predictive performance of convolutional neural networks on the cross-validation dataset across multiple encoding methods

We evaluated the predictive performance of thirteen amino acid encoding methods using PEPIC on the cross-validation dataset (Fig. S1). Performance was assessed on the validation sets using micro-averaged F1-score across twenty repetitions to ensure statistical robustness. Among the tested encodings, Miyazawa energies achieved the highest micro-averaged F1-score (0.991 ± 0.007), followed closely by hydrophobicity matrix (0.990 ± 0.005), binary 5-bit encoding (0.989 ± 0.010), and BLOSUM62 (0.989 ± 0.006). Other encoding methods, such as one-hot (0.988 ± 0.008), AESNN3 (0.988 ± 0.011), and PAM250 (0.984 ± 0.014), also demonstrated competitive performance. Despite slight variations in the results, none of the differences were statistically significant. Given its strong performance, we adopted the Miyazawa energies encoding for the remainder of this study.

**Supplementary Table 1. Sequences for interpretation of plastics-hydrolyzing enzymes.**

| Class | Accession number (Nr) | Sequence |
|---|---|---|
| PET | MAA58622.1 | MNTYLLRTLSICLFAGLFSMQVQAITPDPEPDPDPDPPSTCSNCYQRGPNPTVSALEADSGPYSVRTINVSSWVSGFGGGTIHYPVGTEGTMGAIAVIPGYVSYERSIKWWGPRLASWGFVVITTDTNTIYDQPDSRADQLSAALDYVISQSNSSRSPIYGMVDANRLGAMGWSMGGGGTLKLSTERELKAAIPQAPYYAGFNPFDEITTPTLIIACELDVVAPVAQHASPFYREIPGSTAKAFLEINGGDHFCANSGYPDEDILGKYGIAWMKRFIDEDRRYDQFLCGPNHEADRSISEYRDTCNY |
| | RLP53020.1 | MLEGRYMKTVRFNTAAAVFTSALLSSQVFAITDDPVDPVDPVDPPSSGTVRGPDPTLSALESTRSGPYSVRTENVSNLSASGFGGGTIHYPTNAGENMGAIAVIPGYVSYESSIEWWGPRLASWGFVVITIDTNTIYDQPDSRADQLSAALDHLIDESGSSSSPISGLVDASRLGVIGWSMGGGGTLKLATERNLKAIIPQAPWYSGFNSFDRITTPTMIIACESDAIAPVGQHASPFYNDIPNSTAKAFLEINGGSHYCANSGYSDEDILGKYGISWMKRFMDNDTRYSQFLCGPNHESDRSISEYRDTCNY |
| | WP_021018894.1 | MNVLTKCKLALGIIAIFFSLPSFAVPCSDCSNGFERGQVPRVDQLESSRGPYSVKTINVSRLARGFGGGTIHYSTESGGQQGIIAVVPGYVSLEGSIKWWGPRLASWGFTVITIDTNTIYDQPDSRASQLSAAIDYVIDKGNDRSSPIYGLVDPNRVGVIGWSMGGGGSLKLATDRKIDAVIPQAPWYLGLSRFSSITSPTMIIACQADVVAPVSVHASRFYNQIPGTTPKAYFEIALGSHFCANTGYPSEDILGRNGVAWMKRFIDKDERYTQFLCGQNFDSSLRVSEYRDNCSYY |
| | WP_077315388.1 | MKFLIKVNFILSIFAVFISPQIFAATVACSDCSNGFQRGTLPRVDQLESSRGPYSVKTSNVSVFARGFGGGTIHYSTDSGGQQGIIAVIPGYVSYESSIKWWGPRLASWGFTVITINTNTIYDQPNSRANQLSAAIDYVIDKGNDRSSPIYGLVDPERVGVIGWSMGGGGTLKLATDRDIDAIIPQAPWYSGLNSFSRITTPTMVIACQADAVAPVALHASIFYNQIPRSTPKAFFEIAAGSHFCGNSGYPNEDILGRNGVAWMKRFIDNDTRYNQFLCGQNFDRSLRVSDYKDTCNTY |
| | RMH89651.1 | MKLKAYLARITSLVTVSLLASSLAYAAPGPSAPCADCSRGPNPTVASLQSRSGPFTVSTFSVSGYLRGFGNSTVHYPTNATGKMGAIAVIPGYLSYEDSIRWWGPRLASHGFVVITMNTNTIYDQPDSRATQLSRALDYVIEQSNSRSSPISGKVDSTRLGAIGWSMGGGGSLKLSTERSLNAIIPQAPYYAGLNRFNTINTPTMILACSADVVAPVGSHASPFYNRIPEATPKAFLEIYGGSHFCANSGYPNEDLLGMYGISWMKRFIDFDSRYSQFLCGPNHAADLRISEYRENCNY |
| | OUS39971.1 | MPNPNPAPCQEDCDFTRGPDPTISSLEASAGPYSVANQGVSRSVDGFGGGTIFYPMNTTGTMGAIAIAPGFLAGESSIEWWGPRFASHGFVIITIATNSVFDQPNSRETQLSSALDYVISQSNSGNSPISGMVDSTRVGAMGWSMGGGGALRLASGDRLSAVIPLAPWHQGRNSFDQLETPTLIIACENDTVAPVNRHASSFYNSIPSSTDKALLEISNGAHSCANGGGANGGLLGKYGVSWMKRFIDNDLRYDQFLCGPNHAANSAVSEYRGTCNY |
| | WP_085988667.1 | MPLSMNKLQKLPLTLTSAALLCAGGLTVNTAVAETRGPDPTEAYVEAEGPYNVDTINVSSLALGFGGGTIHYPTNTTGQMGGIVVIPGYLSYESSIEWWGERLASHGFVVMTIDTNTIYDQPGSRRDQIDAALDYLVDRSDSSFSAISGMVDGDRLGAVGWSMGGGGTLQLASGDRLSAAIPLAPWNSSFNDFDDIETPTLIFACENDTVAPVGVHASPFYYDIPASTDKAFFEINNGNHFCANGDNANDAVLSKYGVSWMKLHIDQDARYGQFLCGPNHESQYRISEYRGTCPY |
| | MBF78136.1 | MPFNKKGILAACGAGALLFSMSALANNPPPTDPDPGDGGSSPYQRGPDPSVSFLEADRGNYSVSTSRVSGLVSGFGGGTIHYPSGTTGTMAAIVVIPGFVSAESSIEWWGPKLASYGFVVMTIDTNSGFDQPGSRATQINNALDYLVSQNTSSS |

| | | |
|---|---|---|
| | | SPVRGMIDTSRLGVVGWSMGGGGTLRVAREGRIKAAIPLAPWDTSTYYSS<br>RSQAPTLIFACESDVIAPVYQHASPFYNALPSSIDKAFVEINNGSHYCGNGG<br>SIYNDVLSRFGVSWMKLHLDEDARYKQFLCGPNHTSDSQISDYRGNCPY |
| | WP_1181318<br>88.1 | MSALANNPPPTDPDPGNGGSSPYQRGPDPSVNFLEADRGQYNVDDERVSSF<br>VSGFGGGTIHYPTGTTGTMAAIVVIPGFVSAESSIEWWGPKLASYGFVVMTI<br>DTNSGFDQPGSRATQINNALDYLVDQNTSVGSPVRGMIDTDRLGVIGWSM<br>GGGGTLRVGREGRIKAAIPLAPWDTSSYYASRAQAPTLIFACESDVIAPVYQ<br>HASPFYNALPSNIDKAFVEINNGSHYCGNGGSIYNDVLSRFGVSWMKLHLD<br>EDARYKQFLCGPNHTSDSQISDYRGNCPY |
| | WP_0918498<br>96.1 | MSDHYASNPLRSVVAAASLLFSASVFAAGGGGSDGGDDGCTSNCGYERGP<br>APTESFLEASSGPYSVRTDRVSSLVGGFGGGTIHYPTGTSGTMGAVVVIPGF<br>VSAESSIEWWGPKLASHGFVVMTIDTNSGFDQPPSRATQINNALDYLIEQN<br>GSSSSPVSGMIDTNRLGVIGWSMGGGGTLRVASEGRIQAAIPLAPWDTSSFR<br>FRNIETPTLIIACESDIIAPVRSHADPFYEAIPSSTDKAFVELNNGSHYCGNGG<br>NSYNDVLSRFGVSWMKLHLDNDQRYNQFLCGPDHERDWDISEYRGTCPY |
| | CAH17554.1 | MAVMTPRRERSSLLSRALRFTAAAATALVTAVSLAAPAHAANPYERGPNP<br>TDALLEARSGPFSVSEERASRFGADGFGGGTIYYPRENNTYGAVAISPGYTG<br>TQASVAWLGKRIASHGFVVITIDTNTTLDQPDSRARQLNAALDYMINDASS<br>AVRSRIDSSRLAVMGHSMGGGGSLRLASQRPDLKAAIPLTPWHLNKNWSS<br>VRVPTLIIGADLDTIAPVLTHARPFYNSLPTSISKAYLELDGATHFAPNIPNKI<br>IGKYSVAWLKRFVDNDTRYTQFLCPGPRDGLFGEVEEYRSTCPF |
| | G8GER6.1 | MPPHAARPGPAQNRRGRAMAVITPRRERSSLLSRALRFTAAAATALVTAVS<br>LAAPAHAANPYERGPNPTDALLEARSGPFSVSEERASRFGADGFGGGTIYY<br>PRENNTYGAVAISPGYTGTQASVAWLGERIASHGFVVITIDTNTTLDQPDSR<br>ARQLNAALDYMINDASSAVRSRIDSSRLAVMGHSMGGGGTLRLASQRPDL<br>KAAIPLTPWHLNKNWSSVRVPTLIIGADLDTIAPVLTHARPFYNSLPTSISKA<br>YLELDGATHFAPNIPNKIIGKYSVAWLKRFVDNDTRYTQFLCPGPRDGLFG<br>EVEEYRSTCPF |
| | WP_1247733<br>20.1 | MSSPTTTRPRSVVARLALAAVLAAGGVLAGPAGVAQAASPYERGPAPTTAI<br>LEASRGPFATASQSVSSLSVVGFGGGVIYYPTSTSEGTFGAIAISPGFTASWS<br>SISWLGPRIASHGFVVIGIETNTRLDQPDSRGRQLLAALDYLTERSSVRSRID<br>SSRLAVAGHSMGGGGSLEAASSRPSLQAAVPLAPWNTDKSWSELRVPTLII<br>GGESDSVAPVATHSVPFYNSIPASAEKAYLELNGASHFFPQTTNTPTARQM<br>VAWLKRFVDDDTRYEQFLCPGPSGSQIQEYRNTCPSA |
| | TKK88911.1 | MSRIATAALAFTLATGTAVTLAPSAQAAGFERGPNPTSAILEASRGPFSVAT<br>TSVSSLVSGFGGGTIYYPTDTSQGTFGAIAISPGYTARWSSLEWLGPRIASHG<br>FVVIGIETNSTLDQPASRGNQLLAALDYLVNSSSTTVRSRIDRNRLAVAGHS<br>MGGGGTLHAAEDRPSLKAAVPIAPWNTDKTWGSVRVPTLIVAGESDSVAS<br>PTTHASPFYNSITQTEKAYLELNSASHFFPQTTNTPFAKQFVAWLKRWVDE<br>DTRYSQFICPGPSGLAIEEYRSTCPV |
| | WP_1608756<br>56.1 | MTTTTWRTRIASLALAAAAATGLTSGVGLGAPISAVAATANPYERGPAPTR<br>ASIEATRGAYATAQTSVSSLVSGFGGGTIYYPTTTADGTFGAVIIAPGYTAT<br>SSSLAWLGPRLASQGFVVFTIDTDSRYDQPASRGDQLLEAADYLTRTSVVA<br>SRVDARRVALMGHSMGGGGTLEAIKDRPSIKAAIPLTPWNLDKTWPEVTT<br>PTLIIGADNDSVAPVASHAEPFYGSLPSTLDKAYLELRNASHFAPNSANTTI<br>ASYSIAWLKRFVDDDTRYSQFLCPTPASSLAIAEYRSTCPY |
| | WP_1425690<br>56.1 | MVTRLALVVLLALAGGLLTAPAAHAAVHGPDPTDALLESSRGPYATAQTD<br>VSSLSVSGFGGGTIYYPTTTSEGTFGGVAIAPGYTADKSSLAWLAARLASH<br>GFVVFNIDTLTRLDQPDSRGRQLLAALDYLTQRSSVRGRVDATRLGVMGH<br>SMGGGGTLEAVDDRPSVRAAVPLTPWNLDKTWSGVRTPTLIIGAEADTVA<br>PVASHAVPFYTSLSSSLDKAYLELNGATHFAPNTTNTTIGKYAVAWMKRF<br>VDDDTRYDQFLCPGPGRSLTVEEYRSTCPF |

| | | |
|---|---|---|
| | EFL43114.1 | MHRPAGGSPRQRGPLVVQQHPHTGGRRTGRFAALAAAVAAVVGLTTLGG<br>PGAHAADNPYERGPAPTESSIEALRGPYAVSETSVSSLSVTGFGGGTIYYPT<br>STADGTFGAIAVSPGFTAYQSSIAWLGPRLASQGFVVFTIDTNTTLDQPASR<br>GDQLLAALDYLTQRSAVRGRIDSSRLGVMGHSMGGGGTLEAAKDRPSLQ<br>AAIPLTPWNLDKTWPEVRTPTLLFGADGDTVAPVGTHAEPLYTGLPSSLDR<br>AYLELNGATHFTPNSSNTTIAKYSISWLKRFIDNDTRYEQFLCPLPRPSLTIE<br>ESRGNCPHTS |
| | WP_0855761<br>51.1 | MQQHPHTSRRGTGRFAALTAAVAAVVGLTTLNGPGAQAADNPYERGPAP<br>TESSIEALRGPYAVSDVSVSSLSVTGFGGGTIYYPTSTADGTFGAIAISPGFT<br>AYQSSIAWLGPRLASQGFVVFTIDTNTTLDQPASRGDQLLAALDYLTQRSA<br>VRGRVDSSRLGVMGHSMGGGGTLEAAKDRPSLQAAIPLTPWNLDKTWPE<br>VRTPTLLFGADGDTVAPVSSHAEPLYSGLPSSLDRAYIELNGATHFTPNSSN<br>TTIAKYSVSWLKRFIDNDTRYEQFLCPLPRPSLTVEESRGNCPHTS |
| | WP_1504740<br>04.1 | MQQQARTGAAHRVPTGSPHRRSARFAGLATAIAAVVGLTTLNGAGAQAA<br>DNPYERGPAPTTSSIEAARGSYSVSQTSVSSLAVTGFGGGTIYYPTSTADGT<br>FGAVAISPGYTGTQSTMAWLGPRLASQGFVVFTIDTNTTLDQPDSRGRQLL<br>AALDYLTQRSSVRGRVDSTRLGVMGHSMGGGGSLEAAKTRPSLQAAIPLT<br>PWNTDKSWPEISTPTLIFGADGDTIAPVASHAEPFYSSLPSSLDRAYLELNGT<br>SHLTPISSNTTIAKYSVSWLKRFIDNDTRYEQFLCPLPRPSLTIEEYRGNCPHT<br>S |
| | WP_0537570<br>25.1 | MQQHPRSTTASAAPGPARGAGRRGTRRFAGAAAAIAAAVALSTLTGPGAR<br>AADNPYERGPAPTTASIEASRGPYSVSETSVSSLAVSGFGGGTIYYPTSTAD<br>GTFGAVAVSPGYTGTQSSIAWLGPRLASQGFVVFTIDTLTTLDQPDSRGRQ<br>LLAALDYLTTRSSVRGRVDSTRLGVMGHSMGGGGSLEAAKSRPSLQAAIP<br>LTPWNLDKSWPEVTTPTLIVGADGDSIAPVSSHAEPFYGSLRSSLDRAYLEL<br>NGASHFTPNSSNTTIAKYSVSWLKRFIDNDTRYEQFLCPLPSPSLTIEEYRGN<br>CPHTS |
| | BCL25765.1 | MQQHSQNSSLTAGTVPPESAGRRPRRRDGRGGWAAKRITGALAALTTVVG<br>LSSLASPGAHAADNPYERGPAPTTSSIEAARGSYSVSQTTVSSLAVTGFGGG<br>TVYYPTSTADGTFGAVVISPGYTGTQSSISWLGARLASQGFAVFTIDTLTTL<br>DQPDSRGRQLLAALDYLTERSSVRTRVDGSRLAVMGHSMGGGGSLEAAK<br>SRPSLQAAIPLTPWNTDKSWPEVSTPTLIVGADGDTIAPVASHAEPFYGSLPS<br>STDKAYLELNNATHFTPNSSDTTIAKYSISWLKRFVDNDTRYEQFLCPLPRP<br>SLTIEEYRGNCPHTS |
| | WP_0699337<br>76.1 | MRNAPAHRRRGSGRLRSLVAGLAALLAVGGLSSVATPAAQAADNPYERG<br>PAPTTASIEAPNGPYAVSQTSVSSLSVTGFGGGTVYYPTTTGDGTFGAVAIS<br>PGFTAGESSIAWLGPRLASQGFVVFTIGTLTRYDQPDSRGSQLLAALDYLTQ<br>RSTVRARIDSGRLGVMGHSMGGGGSLEAAKSRPSLQAAIPLTGWNTDKTW<br>PEIKTPTLVVGADGDTVASVGSHSEPFYESLPSSLDKAYLELNNATHFTPNT<br>SNTTIAKYSISWLKRFIDNDTRYEQFLCPLPRPSLTIEEYRGNCPHTS |
| | WP_2373253<br>67.1 | MQQHLLARRQTPHPSRSRTLTGLLTAAAATAGLLLTALAPGAQAVAANPY<br>ERGPAPTNASIEASRGSYATSQTSVSSLAVSGFGGGTIYYPTSTADGTFGAV<br>VISPGFTAYQSSIAWLGPRLASQGFVVFTIDTNTTLDQPDSRGRQLLSALDY<br>LTQRSSVRTRVDASRLGVMGHSMGGGGSLEAAKSRTSLKAAIPLTGWNTD<br>KTWPELRTPTLVVGADGDTVAPVGTHSKPFYESLPGSLDKAYLELRGASHF<br>TPNSSDTTIAKYSLSWLKRFIDNDTRYEQFLCPIPRPSLTIAEYRGTCPHTS |
| | WP_1612676<br>37.1 | MQQHLLARRQAPRPSRSRTLTGLLTAAAATAGLLLTGLAPGAQAVAANPY<br>ERGPAPTNASIEASRGSYATSQTSVSSLAVSGFGGGTIYYPTSTADGTFGAV<br>VISPGFTAYQSSIAWLGPRLASQGFVVFTIDTNTTLDQPDSRGRQLLSALDY<br>LTQRSSVRTRVDASRLGVMGHSMGGGGSLEAAKSRTSLKAAIPLTGWNTD<br>KTWPELRTPTLVVGADGDTVAPVATHSKPFYESLPGSLDKAYLELRGASHF<br>TPNSSDTTIAKYSLSWLKRFIDNDTRYEQFLCPIPRPSLTIAEYRGTCPHSS |

| | | |
|---|---|---|
| | AOS64284.1 | MQSSSIASRRARVRSAGRPRTRLAGLVLALTMVATGLAAAPAATAQENPY<br>ERGPAPTERSIEALRGPFAVAEDRVSSLVIGFGGGTIYYPTDTSEGTFGAVA<br>VSPGYTGTQSSMAWLGPRLASQGFVVFTIDTNTTVDQPDSRGRQLLAALD<br>YLVEDSDVRNRIDPNRLGVMGHSMGGGGSLSAAESRPALQAAIPLTGWHL<br>SKNWSRVTVPTLVVGAENDLIAPVRSHSIPFYESLSSSLDKAYLELDGASHF<br>APNISNTTIAKYSISWLKRFIDDDLRYEQFLCPPPDDREISEYRNTCPHS |
| | KOV83888.1 | MPNEVYSAVQLRTLIPLALTLVLVTGTAAQAADNPYERGPAPTVSSIEALR<br>GPFAVSETSVSSLVGGFGGGTIYYPTSTTSGTFGAVAVSPGYTGTQSSISWL<br>GPRLASQGFVVFTIDTNTIYDQPDSRASQLLAALDYLTQQSSVRSRIDATRL<br>GVMGHSMGGGGTLRAASQRPTLQAAIPLTAWHTTKNWSSVRVPTLVVGA<br>EDDSIAPVATHSEPFYTTLPSTLDKAYLELNNATHFAPNSNNTTIAKYSISW<br>LKRFIDNDTRYEQFLCPAPGRSTLIEEYRDTCPHS |
| | WP_373873195.1 | MDKVIPKLFGIAAAVALGGAGITLIPDADAATASFAKGPAPSNASIEAVRGP<br>FAVAQSNVSRASVSGFGGGDIYAPTDTGAGTFGAVVIAPGFTARKSSMAW<br>LAPRLASQGFVVFNIDTLSTSDQPASRGRQLLAAADFLTQRSTVRARIDAGR<br>VAVIGHSMGGGGALEAAGSRPALAAAIPLTPWNLTKSFSRNAVPTLVIGAE<br>ADSIAPVRSHAQPFFQSLPAVPGKAFLNLNGASHFAPNTPNTTIAKFSISWL<br>KLFVDDDTRYQQFVCPGPGAGAAVQEYRSTCDQFTP |
| | WP_328594753.1 | MPRTTLRTLAAAVLAAGAVGVLPTAPAHAAGFERGPAPTEASVTAAKGPF<br>AIDRIEVPAGSGTGFNSGTIYYPTSTAEGTFGAVAISPGFVSPKSWIDWYGPR<br>LASQGFVVMTLETFSYFDAPDGRADQLLAALDYLTAKSKVKDRIDPNRLA<br>AMGHSMGGGGALSAAVKRPSLKAVVPLAPWYVGGGLEQSTVPTMIFGAD<br>NDFIAPVASNARPFYQSLTKVPEKAYLELENAGHVGSFNSPNTTIAKYAISW<br>LKRFVDDDTRYSQFLCPAPKFPSSTIQEYRDTCPS |
| | G9BY57.1 | MDGVLWRVRTAALMAALLALAAWALVWASPSVEAQSNPYQRGPNPTRS<br>ALTADGPFSVATYTVSRLSVSGFGGGVIYYPTGTSLTFGGIAMSPGYTADAS<br>SLAWLGRRLASHGFVVLVINTNSRFDYPDSRASQLSAALNYLRTSSPSAVR<br>ARLDANRLAVAGHSMGGGGTLRIAEQNPSLKAAVPLTPWHTDKTFNTSVP<br>VLIVGAEADTVAPVSQHAIPFYQNLPSTTPKVYVELDNASHFAPNSNNAAIS<br>VYTISWMKLWVDNDTRYRQFLCNVNDPALSDFRTNNRHCQ |
| | P19833.1 | MFIMIKKSELAKAIIVTGALVFSIPTLAEVTLSETTVSSIKSEATVSSTKKALP<br>ATPSDCIADSKITAVALSDTRDNGPFSIRTKRISRQSAKGFGGGTIHYPTNAS<br>GCGLLGAIAVVPGYVSYENSIKWWGPRLASWGFVVITINTNSIYDDPDSRA<br>AQLNAALDNMIADDTVGSMIDPKRLGAIGWSMGGGGALKLATERSTVRAI<br>MPLAPYHDKSYGEVKTPTLVIACEDDRIAETKKYANAFYKNAIGPKMKVE<br>VNNGSHFCPSYRFNEILLSKPGIAWMQRYINNDTRFDKFLCANENYSKSPRI<br>SAYDYKDCP |
| PCL | WP_025804184.1 | MSTLSWVRGVNGTLGWVAPKLVASKMRLAFMTPREHLPRDWELPLLARS<br>ERITLRFGLSALRWGQGPAVLLMHGWEGRPTQFASLIDALVGAGYSVVAL<br>DGPAHGRSPGHEANVMLFARAMLEAAAELPPLRAVIGHSMGGASAMLAV<br>QLGLRTETLVSIAAPARILGVLRGFARYVRLPPKARSVFIRQVEQDVGMRA<br>AAMDVAHYQLDMPGLIVHAEDDNFVPVKESELIHDAWFDSRLLRLKEGG<br>HQRVLADPRVIEGVLTLLAGRSLQARQSA |
| | WP_059181212.1 | MSTLKWVRGVNGTLGWFAPQLVASRMRLAFMSPRALPPRDWELPLLAKS<br>ERITLRFGLSALRWGQGPAVLLMHGWEGRPTQFASIITALVDAGYSVVALD<br>GPAHGRSPGEEANVVLFARAMLEAAAELPPLQAVIGHSMGGASAMLAVQ<br>LGLRTETLVTIAAPARILGVLRGFAKYVRLPPKARSAFIRQVEKDVGMRAA<br>ALDVAHYQLDMPGLIVHAEDDRFVSAKESQLIHEAWFDSRLLRLEEGGHQ<br>RVLADPRVIDGVLSLLAGRSLHSRQSA |
| | WP_052145136.1 | MERLYRGMGRAALDRAYNNTRAIANFPAVLADFRTRSAALYERVRGRRD<br>LRYGDRPRERFDWLPGGRANAPTFVFIHGGYWQNCAKEDFAFVAHGPLAR |

| | | |
|---|---|---|
| | | GFNVVLAEYTLAPDASMTQIVDEIGRLIDHLRADRDGLGTAGRPLCLSGHSAGGHLSALHRGHAFVTSALAISPLVDLEPISLSWLNEKLQLSEREIAAYSPLWHVGKGAPTVVAVGADELPELVRQADDYTAACAAAGEPVWGAHVPGCTHFSVLDDLAQPNGTLMRLLDAAIAGSGHGGDERADEERR |
| | WP_124320071.1 | MSTLSWVRGVNGTLGWVAPKLVASKMRLAFMTPRERLPRDWELPLLARSERITLRFGLSALRWGQGPAVLLMHGWEGRPTQFASLIDALVGAGYSVVALDGPAHGRSPGQEANVMLFARAMLEAAAELPPLRAVIGHSMGGASAMLAVQLGLRTETLVSIAAPARILGVLRGFARYVRLPPKARSAFIRQVEQDVGMRAAAMDVAHYQLDMPGLIVHAEDDNFVPVKESDLIHEAWFDSRLLRLKEGGHQRVLADPRVIEGVLTLLAGRSLQARQSA |
| | WP_134174323.1 | MNTLKWIRGVNGTLGWIAPKRVASKMRLAFMTPRSLPLRDWELPLLASSERITLRFGLSALRWGQGPAVLLMHGWEGRPTQFAALITALVDAGYTVVALDGPAHGRSPGREANVVLFARAMLEAAAELPPLQAVIGHSMGGASAMLAVQLGLRTETLVSIAAPARILGVLRGFARYVGMPPRARSAFIRQVEQDVGMRAATLDVAHYQLDMPGLIVHAEDDNFVSVKESQLIHESWFDSRLLRLEGGGHQRVLADPRVIDGVLSLLAGRSLQARQSA |
| | WP_096796149.1 | MNTLKWVRGVNGTLGWIAPQRVASKMRQAFMTPRTLPLRDWELPLLASAERITLRFGLSALRWGQGPAVLLMHGWEGRPTQFAALITALVEAGYTVVALDGPAHGRSPGREANVVLFARAMLEAAAELPPLQAVIGHSMGGASAMLAVQLGLRTETLVSIAAPARILGVLRGFARYVGMPPRARSAFIRQVEQDVGMRAATLDVAHYQLDMPGLIVHAEDDTFVSVKESQLIHESWFDSRLLRLESGGHQRVLADPRVVDGVLSLLAGRSLQARQSA |
| | WP_054596612.1 | MSTLSWVRGVNGTLGWVAPQWVASKMRSVFMTPRELPPRDWEMPLLAKSERITLRFGLSALRWGQGPTVLLMHGWEGRPTQFASLITALVDAGYTVVALDGPAHGRSPGREANVVLFARAMLEAAAELPPLQAVIGHSMGGASAMLAVQLGLRTETLVSIAAPARILAVLRGFARHVRMPPKVRSAFIRKVERDVGIQASRLDVAHYQLDMPGLIVHAEDDVFVSVNESQLIHDAWFDSRLLRLEEGGHQRVLADPRVIEGVLSLLSGRSLQARQSA |
| | WP_123592831.1 | MNTLKWVRGVNGTLGWIAPQRVASKMRLAFMTPRSLPLRDWELPLLASSERITLRFGLSALRWGQGPTVLLMHGWEGRPTQFAALITALVDAGYTVVALDGPAHGRSLGREANVVLFARAMLEAAAELPPLQAVIGHSMGGASAMLAVQLGLRTETLVSIAAPARILGVLRGFARYVGMPPRARSAFIRQVEQDVGMRAATLDVAHYQLDMPGLIVHAEDDNFVSVKESQLIHESWFDSRLLRLEGGGHQRVLADPRVVDGVLSLLAGRSLQARQSA |
| | WP_059728396.1 | MTILYRGMDRAALDAAYLNTKVVPDFPALLASMQARSAALYDTAHGRRDLRYGAQPAQRFDWLSCGRPDAPLFVFIHGGYWQHCAKEDFAYAASGPLARGFDVILAEYTLAPVATMTGIVGEIGALLDYLANDPDAIGTAGRPIHLSGHSAGGHLTAVYRAHPAVVSALAISPLVDLEPISLCVLNDKLQLGAREVDAYSPLRHVGPGAPTVVAVGDAELPELVRQARDYATACEAAGERVVHVGLPGMRHFDVLDDLAKPDGAMLAALQSIAPR |
| | WP_038631269.1 | MSTLSWVRGVNGTLGWVAPTLVASKMRLAFMTPRERLPRDWELPLLARSERITLRFGLSALRWGQGPAVLLMHGWEGRPTQFASLIDALVGAGYSVVALDGPAHGRSPGHEANVMLFARAMLEAAAELPPLRAVIGHSMGGASAMLAVQLGLRTETLVSIAAPARILGVLRGFARYVRLPPKARSAFIRQVEQDVGMRAAAMDVAHYQLDMPGLIVHAEDDNFVPVKESDLIHEAWFDSRLLRLKEGGHQRVLADPRVIEGVLTLLAGRSLQARQSA |
| | AIC18917.1 | MTMTLLYRDMNQAQLDAAYNNTQAVPDFPGIYAALQARSASFYASAAGRLNLPYGTAPRQRYDWLPCGKADAPTLIFIHGGYWQNCSKEDFAFIAAGPLAAGFNIVLAEYTLAPQASMTQIVSEIGSLLEHLQADADQLGIAGHKVVLSGHSAGGHLALQFRSHPWVTDVLAISALVDLEPISLSWLNEKLSLSEAEIDAYSPLYHIDKGANTWVAVGADELSELVRQSDEYAKQALARGESVQLIHVPGCTHFSVLDEMAKPQGALLQALSSIR |

| | | |
|---|---|---|
| | WP_139055557.1 | MNTLKWVRGVNGTLGWIAPQRVASKMRQAFMTPRTLPLRDWELPLLASSERITLRFGLSALRWGQGPAVLLMHGWEGRPTQFAALITALVEAGYTVVALDGPAHGRSPGREANVVLFARAMLEAAAELPPLQAVIGHSMGGASAMLAVQLGLRTETLVSIAAPARILGVLRGFARYVGMPPRARSAFIRQVEQDVGMRAATLDVAHYQLDMPGLIVHAEDDTFVSVKESQLIHESWFDSRLLRLESGGHQRVLADPRVVDGVLSLLAGRSLQARQSA |
| | WP_103304852.1 | MNTLKWIRGVNGTLGWIAPKQVASKMRLAFMTPRALPLRDWELPLLANSERITLRFGLSALRWGQGPTVLLMHGWEGRPTQFAALITALVEAGYTVVALDGPAHGRSPGREANVVLFARAMLEAAAELPPLQAVIGHSMGGASAMLAVQLGLRTETLVSIAAPARILGVLRGFARYVGMPPRARSAFIRQVEQDVGMRAATLDVAHYQLDMPGLIVHAEDDNFVSVKESQLIHESWFDSRLLRLESGGHQRVLADPRVIDGVLSLLAGRSLQARQSA |
| | WP_007969226.1 | MNTLKWVRGVNGTLGWIAPKQVASKMRTAFMTPRALPLRDWELPLLASSERITLRFGLSALRWGQGPAVLLMHGWEGRPTQFAALITALVEAGYTVVALDGPAHGRSPGREANVVLFARAMLEAAAELPPLQAVIGHSMGGASAMLAVQLGLRTETLVSIAAPARILGVLRGFARYVGMPPRARSAFIRQVEQDVGMRAATLDVAHYQLDMPGLIVHAEDDTFVSVKESQLIHESWFDSRLMRLESGGHQRVLADPRVVDGVLSLLAGRSLQARQSA |
| | WP_047600195.1 | MNTLKWVRGVNGTLGWIAPQRVANKMRQAFMTPRTLPLRDWELPLLASSERITLRFGLSALRWGQGPAVLLMHGWEGRPTQFAALITALVEAGYTVVALDGPAHGRSPGREANVVLFARAMLEAAAELPPLQAVIGHSMGGASAMLAVQLGLRTETLVSIAAPARILGVLRGFARYVGMPPRARSAFIRQVEQDVGMRAATLDVAHYQLDMPGLIVHAEDDNFVSVKESQLIHESWFDSRLLRLESGGHQRVLADPRVVDGVLSLLAGRSLQARQSA |
| | WP_130927896.1 | MNALKWVRGVNGTLGWFAPKLVASKMRLAFMTPRALPLRDWELPLLASSERITLRFGLSALRWGQGPTVLLMHGWEGRPTQFASLINALVDAGYTVVALDGPAHGRSPGREANVVLFARAMLEASAELPPLQAVIGHSMGGASAMLAVQLGLRTETLVSIAAPSRILGVLRGFARMVGMPPRARSAFIRQVEQDVGMRAATLDVAHYQLDMPGLIVHAEDDNFVSVKESQLIHEAWFDSRLLRLESGGHQRVLADPRVIDGVLSLLAGRSLQARQSA |
| | WP_123463437.1 | MNTLKWIRGVNGTLGWIAPKRVASKMRLAFMTPRTLPLRDWELPLLANSERITLRFGLSALRWGQGPTVLLMHGWEGRPTQFAALITALVEAGYTVVALDGPAHGRSPGREANVVLFARAMLEAAAELPPLQAVIGHSMGGASAMLAVQLGLRTETLVSIAAPARILGVLRGFARYVGLPPRARSAFIRQVEQDVGMRAATLDVAHYQLDMPGLIVHAEDDTFVSVKESQLIHESWFDSRLLRLESGGHQRVLADPRVVDGVLSLLAGRSLQARQSA |
| | WP_151551651.1 | MNALKWVRGVNGTLGWFAPKLVASKMRLAFMTPRALPLRDWELPLLASSERITLRFGLSALRWGQGPTVLLMHGWEGRPTQFASLINALVDAGYTVVALDGPAHGRSPGREANVVLFARAMLEAAAELPPLQAVIGHSMGGASAMLAVQLGLRTETLVSIAAPARILGVLRGFARMVGMPPRARSAFIRQVEQDVGMRAATLDVAHYQLDMPGLIVHAEDDNFVSVKESQLIHEAWFDSRLLRLESGGHQRVLADPRVIDGVLSLLAGRSLQARKSA |
| | WP_123448644.1 | MNTLKWVRGVNGTLGWIAPKRVASKMRLAFMTPRSLPLRDWELPLLASSERITLRFGLSALRWGQGPTVLLMHGWEGRPTQFAALITALVDAGYTVVALDGPAHGRSPGREANVVLFARAMLEAAAELPPLQAVIGHSMGGASAMLAVQLGLRTETLVSIAAPARILGVLRGFARYVGMPPRARSAFIRQVEQDVGMRAATLDVAHYQLDMPGLIVHAEDDSFVSVKESQLIHEAWFDSRLLRLEGGGHQRVLADPRVIDGVLSLLAGRSLQARQSA |
| | WP_127650388.1 | MNTLKWVRGVNGTLGWIAPKRVASKMRLAFMTPRALPLRDWELPLLASSERITLRFGLSALRWGQGPAVLLMHGWEGRPTQFAALITALVEAGYTVVALDGPAHGRSPGREANVVLFARAMLEAAAELPPLQAVIGHSMGGASAMLAVQLGLRTETLVSIAAPARILGVLRGFARYVGMPPRARSAFIRQVEQDVGMRA |

| | | |
|---|---|---|
| | | ATLDVAHYQLDMPGLIVHAEDDTFVSVKESQLIHESWFDSRLLRLESGGHQ<br>RVLADPRVVDGVLSLLAGRSLQARQSA |
| | WP_160057137.1 | MNTLKWVRGVNGTLGWIAPQRVASKMRLAFMTPRALPLRDWELPLLASS<br>ERITLRFGLSALRWGQGPAVLLMHGWEGRPTQFAALITALVDAGYTVVAL<br>DGPAHGRSPGREANVVLFARAMLEAAAELPPLQAVIGHSMGGASAMLAV<br>QLGLRTETLVSIAAPARILGVLRGFARYVGMPPRARSAFIRQVEQDVGMRA<br>ATLDVAHYQLDMPGLIVHAEDDNFVSVKESQLIHESWFDSRLLRLEGGGH<br>QRVLADPRVVDGVLSLLAGRSLQARQSA |
| | WP_108591690.1 | MNTLKWVRGVNGTLGWIAPKQVASKMRTAFMTPRSLPLRDWELPLLASS<br>ERITLRFGLSALRWGQGPAVLLMHGWEGRPTQFAALITALVEAGYTVVAL<br>DGPAHGRSPGREANVVLFARAMLEAAAELPPLQAVIGHSMGGASAMLAV<br>QLGLRTETLVSIAAPARILGVLRGFARYVGMPPRARSAFIRQVEQDVGMRA<br>ATLDVAHYQLDMPGLIVHAEDDNFVSVKESQLIHESWFDSRLLRLESGGH<br>QRVLADPRVVDGVLSLLAGRSLQARQSA |
| | WP_332373221.1 | MTILYRGMDRTALDAAYLNTKAVPDFPALLASCQTRSAALYAATPGRRDL<br>RYGAQPAQRFDWLPCGQPDAPLFVFIHGGYWQHCTKEDFAYAASGPLAR<br>GYDVVLAEYTLAPVATMTDIVGEIGALLDHLAADRDGLGTAGRPIHLSGHS<br>AGGHLTAMHRAHPAVVSALAISPLVDLEPIALCCLNDKLQLTAREVDAYSP<br>LHHIGPGAPTVIAVGDAELPELIRQADVYATACEAAGERIARTWLRGMQHF<br>AVLDDLATPDGAMLDALHAIAPR |
| | WP_034186871.1 | MTILYRGMDRAALDAAYLNTKAVPDFPALLASCQARSAALYDETPGRRDL<br>RYGAQPAQRFDWLSCGQAGAPLFVFIHGGYWQHCTKADFAYAASGPLAC<br>GFDVILAEYTLAPVATMTGIVAEIGMLLDHLAADPDRLGTARRPIHLSGHS<br>AGGHLTAMHRAHPAVVSALAISPLVDLEPISLCCLNDKLQLTAHEVDAYSP<br>LRHVGPGAPTVVAVGDAELPELIRQADEYATACEAAGERIARVWLPGMQH<br>FAVLDDLARPDGAMLAALRSITPR |
| | WP_130207001.1 | MSALKWVRGVNGTLGWFAPKLVARKMRLAFMTPRDLPPRDWELPLLAKS<br>ERITLRFGLSALRWGQGPAVLLMHGWEGRPTQFASLITALVDAGYTVVAL<br>DGPAHGRSPGTEANVALFARAMLEAAAELPPLQAVIGHSMGGASAMLAV<br>QLGLRTETLVTIAAPARILGVLRGFARYVGLPPKARSAFIRQVEKDVGMRA<br>ATLDVAHYQLDMPGLIVHAEDDKLVSVKESQAIHEAWFDSRLLRLQEGGH<br>QRVLADPQVIDGVLSLLAGRSLQSRQSA |
| | WP_150773989.1 | MNTLKWIRGVNGTLGWIAPKRVASKMRLAFMTPRALPLRDWELPLLASSE<br>RITLRFGLSALRWGQGPAVLLMHGWEGRPTQFAALITALVDAGYTVVALD<br>GPAHGRSPGREANVVLFARAMLEAAAELPPLQAVIGHSMGGASAMLAVQ<br>LGLRTETLVSIAAPARILGVLRGFARYVGMPPRARSAFIRQVEQDVGMRAA<br>TLDVAHYQLDMPGLIVHAEDDNFVSVKESQLIHESWFDSRLLRLEGGGHQ<br>RVLADPRVVDGVLSLLAGRSLQARQSA |
| | WP_069745339.1 | MTILYRGMDRAALDAAYLNTKAVPDFPALLASCQSRSAALYDAIAGRREL<br>RYGALPAQRYDWLPCGQPGAPLFVFIHGGYWQHCAKEDFAYAASGPLAR<br>GYDVVLAEYTLAPTASMTDIVAEIGALLDHLAADRDGLGIAGRPIHLSGHS<br>AGGHLTAMYRAHPAVAAALSISPLVDLEPISLCCLNDKLQLTAQEIEACSPL<br>RHIGPGAPTVVAVGDAELPELIRQAHDYAAACDAAGERIAHVQLPGMKHF<br>AVLDDLANPDGKMLAALRAIAPR |
| | WP_003226446.1 | MNTLKWVRGVNGTLGWIAPQRVASKMRQAFMTPRSLPLRDWELPLLASA<br>ERITLRFGLSALRWGQGPAVLLMHGWEGRPTQFAALITALVEAGYTVVAL<br>DGPAHGRSPGREANVVLFARAMLEAAAELPPLQAVIGHSMGGASAMLAV<br>QLGLRTETLVSIAAPARILGVLRGFARYVGMPPRARSAFIRQVEQDVGMRA<br>ATLDVAHYQLDMPGLIVHAEDDTFVSVKESQLIHESWFDSRLLRLESGGHQ<br>RVLADPRVVDGVLSLLAGRSLQARQSA |
| | WP_1235329 | MNTLKWVRGVNGTLGWIAPQRVASKMRLAFMTPRVLPLRDWELPLLANS |

| | | |
|---|---|---|
| | 98.1 | ERITLRFGLSALRWGQGPTVLLMHGWEGRPTQFAALITALVDAGYTVVAL<br>DGPAHGRSPGREANVVLFARAMLEAAAELPPLQAVIGHSMGGASAMLAV<br>QLGLRTETLVSIAAPARILGVLRGFARYVGMPRRARSEFIRQVEQDVGMRA<br>ATLDVAHYQLDMPGLIVHAEDDNFVSVKESQLIHESWFDSRLLRLEAGGH<br>QRVLADPRVIDGVLSLLAGRSLQARQSA |
| | WP_1023589<br>40.1 | MSTLKWVRGVNGTLGWFAPKLVASKMRLAFMTPRSLPLRDWELPLLASS<br>ERITLRFGLSALRWGQGPAVLLMHGWEGRPTQFAALITALVEAGYTVVAL<br>DGPAHGRSPGREANVVLFARAMLEAAAELPPLQAVIGHSMGGASAMLAV<br>QLGLRTETLVSIAAPARILGVLRGFARYVGMPPRARSAFIRQVERDVGMRA<br>ATLDVAHYQLDMPGLIVHAEDDSFVSVKESQLIHESWFDSRLLRLESGGHQ<br>RVLADPRVVDGVLSLLAGRSLQARQSA |
| | WP_0079134<br>23.1 | MNTLKWVRGVNGTLGWIAPQRVASKMRLAFMTPRSLPLRDWELPLLASS<br>ERITLRFGLSALRWGQGPTVLLMHGWEGRPTQFAALITALVDAGYTVVAL<br>DGPAHGRSPGREANVVLFARAMLEAAAELPPLQAVIGHSMGGASAMLAV<br>QLGLRTETLVSIAAPARILGVLRGFARYVGMPPRARSAFIRQVEQDVGMRA<br>ATLDVAHYQLDMPGLIVHAEDDNFVSVKESQLIHESWFDSRLLRLEGGGH<br>QRVLADPRVVDGVLSLLAGRSLQARQSA |
| | WP_1507943<br>02.1 | MNTLKWVRGVNGTLGWIAPQRVASKMRLAFMTPRSLPLRDWELPLLASS<br>ERITLRFGLSALRWGQGPTVLLMHGWEGRPTQFAALITALVEAGYTVVAL<br>DGPAHGRSPGREANVVLFARAMLEAAAELPPLQAVIGHSMGGASAMLAV<br>QLGLRTETLVSIAAPARILGVLRGFARYVGMPPRARSAFIRQVEQDVGMRA<br>ATLDVAHYQLDMPGLIVHAEDDNFVSVKESQLIHESWFDSRLLRLEGGGH<br>QRVLADPRVVDGVLSLLAGRSLQARQSA |
| | WP_1332109<br>51.1 | MNTLSWIRSVNGTLGRLAPEHIAGKMRHAFMTPRNLPPRDWELPLLASGE<br>RITLRFGLSALRWGQGPTVLLMHGWEGRPTQFAHLITTLVQAGYTAVALE<br>GPAHGHSPGNQAHVALFARSLLEAAAELPPLRAVIGHSMGGASVMLALQ<br>WGLRAEMAVSVAAPAQLLGVLRNFAHRLGMPSRARAAFVRQVERDVGIPI<br>SRLDVSRYQLEIPALIAHAEDDRIVPASEALTIHQSWFDSRLLLLPEGGHQR<br>VLSDPQLIEGVMALLLRRHSTARQSA |
| | WP_0389941<br>75.1 | MNTLSWIRSVNGTLGHLAPEHVARKMRRAFMTPRNRPPRDWELPLLARAE<br>RITLRFGLSALRWGQGPTVLLMHGWEGRPTQFAHLIDSLVDAGYTAVALE<br>GPAHGHSPGNEANVVLFARALLEAAAELPPLKAVVGHSMGGASMLLALQ<br>WGLRAEVAVSIAAPAQLLGVIRGFARHLGMPARARAAFIRQIERDVGVQIS<br>RLDVSGYQLELPGLIVHAEDDQLVPVDESDAIHRAWFDSRLLRLPDGGHLR<br>VLADPQLREGVLALLQRSSSPARQSA |
| | WP_1600878<br>82.1 | MNSMSWVRGFNASIGLLAPHALASKLRREFMTPHTLPPRDWELPLLAQAE<br>RITLRFGLSALRWGSGPAVLLMHGWEGRPTQFAELIKALVNAGYGVVALD<br>APAHGRSPGREANVVLFARALLEAASELPPLKAVIGHSMGGASALLATQLG<br>LRTEALVSIAAPSRILTMLRRFSHYMGLPRQARAHFVQLVEEQAGIPAGQL<br>DSAHYQLDFPGLVVHAVDDPMVPFSEAEAIHQRWFDSRLLRLERGGHQRV<br>LADPQVVQAVLTLLASLNQAPSNALAS |
| | NMY13981.1 | MNQMAWVRGVNATLGRVAPQLIASRLRERFMTPRTTSPRDWELPLLASSE<br>RITLRFGLSALRWGSGPTVLLMHGWEGRPTQFALLIRGLVDAGYGVVALD<br>APAHGRSPGREANVVLFARALLEAASELPPLRAVIGHSMGGASALLATQM<br>GLRSETLVTIAAPSRILGLLRGFARFMGLPAEARAHFVRQVEKTAGIPAAHL<br>DVQRYRLELPGLIVHAADDQVVPVSEADLIHKAWFDSQLLRLSAGGHQRL<br>LSDPLLLQAVLELLEQVPQASLKALAS |
| PLA | WP_0341262<br>30.1 | MGALTWVRGFNGTVGRLAPHAVASKMRRTFMTPRDLPPRDWELPLLAQS<br>ERITLRFGLSALRWGHGPAVLLMHGWEGRPTQFASLITALVDNGYSVIALD<br>GPAHGRSPGREAHVLLFARAMLEAAAELPPLYAVVGHSMGGASAMLAVQ<br>LGLRTQALVSIAAPSRFLDVLRGFTRMVGLPPRARSAFIQEVELSMGMPLK |

| | | |
|---|---|---|
| | | HLDVAHYHLNIPGLIVHAEDDTFVPVRAAQAIHEAWFDSRLLRLEQGGHQ KVLADPQVIDAVLALLAGCRLQERQSA |
| | WP_032896134.1 | MGALTWVRGFNGTVGRLAPHAVASKMRRTFMTPRDLPPRDWELPLLAQS ERITLRFGLSALRWGQGPAVLLMHGWEGRPTQFASLITALVDNGYSVIALD GPAHGRSPGREAHVLLFARAMLEAAAELPPLYAVVGHSMGGASAMLAVQ LGLRTQALVSIAAPSRFLDVLRGFTRMVGLPPRARSAFIQEVELSMGMPLK HLDVAHYQMNIPGLIVHAEDDTFVPVRAAQAIHEAWFDSRLLRLEQGGHQ KVLADPQVIDAVLALLAGCRLQERQSA |
| | WP_044274707.1 | MGALTWVRGFNGTVGRLAPHAVASKMRRTFMTPRDLPPRDWELPLLAQS ERITLRFGLSALRWGHGPAVLLMHGWEGRPTQFASLITALVDNGYSVIALD GPAHGRSPGREAHVLLFARAMLEAAAELPPLYAVVGHSMGGASAMLAVQ LGLRTQALVSIAAPSRFLDVLRGFTHMVGLPPRARSAFIQEVELSMGMPLK HLDVAHYQMNIPGLIVHAEDDTFVPVRAAQAIHEAWFDSRLLRLEQGGHQ KVLADPQVIDAVLALLAGCRLQERQSA |
| | ABY53108.1 | MGLAIAAAAVFSLPGVATATEPTGGVQPNIVGGGNATQVYSFMVSQQSSS GGHQCGGSLISSTWVVTAKHCGTPYQVRVGTTNRTSGGTVARVAQRIAHP SADLALLRLSTAVPQAPVTIADASGAVGTATRIIGWGQTCAPQGGCGAPITL QELNTSIVSDSRCLGISGASEICTNNPNGNSGACYGDSGGPQIKQVNGVWQ LIGATSRAGNNSSTCATGPSIYVDVPYFRSWIRTNTGV |
| | WP_057005688.1 | MGALTWIRGVNGTLGRLAPHTVANSMRRVFMTPRDLPPRDWELPLLAHA ERVTLRFGLSALRWGQGPAVLLMHGWEGRPTQFASLITALVQNGYSVFAL DGPAHGRSPGREAHVLLFARAMLEAAAELPPLHAVVGHSMGGASAMLAV QLGLRTEALVSIAAPSRFLDVLGGFAGMVGLPSRARSAFIQEVELTLGMPL KHLDVAHYQMDLPGLIVHAEDDTFVPVSASQVIHDAWFDSRLLRLEQGGH QRVLADPRVVEAVLALLAGSCLQARQTA |
| | WP_043206304.1 | MGTLTWIRRFNGTLGRLAPQTVANRMRRAFMTPRDLPPRDWELPLLAQSE RITLRFGLSALRWGQGPAVLLMHGWEGRPTQFASLITALVDNGYSVIALDG PAHGRSPGREAHVLLFARAMLEAAAELPPLHAVIGHSMGGASAMLAVQLG LRTEALVSIASPSRFLDVLRGFTQMVGLPARARSAFIQEVELTFGMPLKHLD VAHYQMNIPGLIVHAEDDTFVPVKASQAIHEAWFDSRLLRLEQGGHQKVL ADPRVIDAVLALLAGRRLQALQSA |
| | WP_116666539.1 | MGALTWVRGFNGTVGRLAPHAVASKMRRTFMTPRDLPPRDWELPLLAQS ERITLRFGLSALRWGHGPAVLLMHGWEGRPTQFASLITALVGDGYSVIALD GPAHGRSPGREAHVLLFARAMLGAAAELPPLYAVVGHSMGGASAMLAVQ LGLRTQALVSIAAPSRFLDVLRGFTRMVGLPPRARSAFIQEVELSMGMPLK HLDVAHYQLNIPGLIVHAEDDTFVPVRAAQAIHEAWFDSRLLRLEQGGHQ KVLADPQVIDAVLALLAGCRLQERQSA |
| | WP_064055161.1 | MMGTLTWIRRFNGTLGRLAPQTVANRMRRAFMTPRDLPPRDWELPLLAQS ERITLRFGLSALRWGQGPAVLLMHGWEGRPTQFASLITALVDNGYSVIALD GPAHGRSPGREAHVLLFARAMLEAAAELPPLHAVIGHSMGGASAMLAVQL GLRTEVLVSIASPSRFLDVLRGFTQMVGLPARARSAFIQEVELTFGMPLKHL DVAHYQMNIPGLIVHAEDDTFVPVKASQAIHEAWFDSRLLRLEQGGHQKV LADPRLIDAVLALLAGRRLQALQSA |
| | WP_076953514.1 | MGTLTWIRRFNGTLGHLAPQTVANKMRRVFMTPRTLPPRDWELPLLAQSE RITLRFGLSALRWGQGPTVLLMHGWEGRPTQFASLITALVDKGYSVIALDG PAHGRSPGREAHVVLFARAMLEAAAELPPLRAVIGHSMGGASAMLAVQL GLRTEALVSIAAPSRCLDALRGFTTMVGLPSRARSAFIREVEMTFAMPLKH LDVAHYQMNIPGLIVHAEDDTFVPVKASQAIHEAWFDSRLLRLEQGGHQK VLADPRVIDATLSLLAGCGLQALQTA |
| | WP_019820855.1 | MGALTWVRGFNGTVGRLAPHAVASKMRRTFMTPRDLPPRDWELPLLAQS ERITLRFGLSALRWGHGPAVLLMHGWEGRPTQFASLITALVDNGYSVIALD |

| | | |
| --- | --- | --- |
| | | GPAHGRSPGREAHVLLFARAMLEAAAELPPLYAVVGHSMGGASAMLAVQ<br>LGLRTQALVSIAAPSRFLDVLRGFTRMVGLPPRARSAFIQEVELSMGMALK<br>HLDVAHYQLNIPGLIVHAEDDTFVPVRAAQAIHEAWFDSRLLRLEQGGHQ<br>KVLADPQVIDAVLALLAGCRLQERQIA |
| | WP_0429370<br>80.1 | MSTLSWIRGINGTLGRVAPRVVASRMRQMFMTPRARLPRDWELPLLATAE<br>RITLRFGLSALRWGKGPTVLLMHGWEGRPTQFANLINALVAAGYTAVALD<br>GPAHGRSPGREANVVVFARALLEAAAELPPLKAVVGHSMGGASAMLATQ<br>LGLRTEALVSIAAPARVLGVLRGFARYVGLPPRARSAFIREVERDVGMRAA<br>HLDIEHYQMDMPGLIVHAEDDRMVRVDESRRIHEAWFDSRLLRLESGGHL<br>QVLADQRLIDGVLALLAGRSLAQRQSA |
| | WP_0995840<br>16.1 | MGALTWIRRFNGTLGHVAPHTVANKMRRAFMTPRKLPPRDWELPLLAQS<br>ERITLRFGLSALRWGQGPAVLLMHGWEGRPTQFASLIKALVDNGYCVIAL<br>DGPAHGRSPGREAHVLLFARAMLEAAAELPPLHAVVGHSMGGASAMLAV<br>QLGLRTQALVSIAAPSRFLDALRGFTRMVGLPARARSAFIQEVEMTFGMPL<br>KYLDVAHYQMNIPGLIVHAEDDTFVPVKASQAIHDAWFDSRLLRLEQGGH<br>QKVLADPRVIEAVLTLLAGCCLQERQSA |
| | WP_0567858<br>24.1 | MGALTWVRGFNGTVGRLAPHAVASKMRRTFMTPRDLPPRDWELPLLAQS<br>ERITLRFGLSALRWGHGPAVLLMHGWEGRPTQFASLITALVDNGYSVIALD<br>GPAHGRSPGREAHVLLFARAMLEAAAELPPLYAVVGHSMGGASAMLAVQ<br>LGLRTQALVSIAAPSRFLDVLRGFTRMVGLPPRARSAFIQEVELSMGMPLK<br>HLDVAHYQLNIPGLIVHAEDDTFVPVRAAQAIHQAWFDSRLLRLEQGGHQ<br>KVLADPQVIDAVLALLAGCRLQERQSA |
| | WP_1976274<br>15.1 | MGALTWIRGFNGTVGRLAPHMVASKLRRTFMTPRNLAPRDWELPLLAQSE<br>RITLRFGLSALRWGQGPAVLLMHGWEGRPTQFASLITALVADGYSVIALDG<br>PAHGRSPGREAHVLLFARAMLEAAAELPPLHAVVGHSMGGASAMLAVQL<br>GLRTEALVSIAAPSRFLDVLRGFAGMVGLPPRARSAFIHEVELTFGMPLKHL<br>DVAHYQMNIPGLIVHAEDDTFVPVNASQAIHDAWFDSRLLRLEQGGHQKV<br>LADPRVIDAVLSLLAGRRLQERQTA |
| | WP_1106820<br>84.1 | MSSMSWIRGFNATVGRLAPDLVASKMHRAFLTPRDLPPRDWELPLLAESE<br>RITLRFGLSALRWGQGPAVLLLHGWEGRPTQFAELIRALVRAGYGVVALD<br>APTHGRSPGHEANVVLFARALLEAAGELPPLKAVIGHSLGGASALLATQLG<br>LRTEALVTIAAPARILGALRRFAHFVGLPKQARARFVRMVEQSAGMPAAQ<br>LDVARYQLDFPGLVVHAEDDPMVPYGEAQSIHAAWPGSRLLPLERGGHSK<br>PLGDPRVVEAVLELLGSADLHSAVSRRVLAATVLAS |
| | XOQ14903.1 | AQSVPWGISRVQAPAAHNRGLTGSGVKVAVLDTGISTHPDLNIRGGASFVP<br>GEPSTQDGNGHGTHVAGTIAALNNSIGVLGVAPSAELYAVKVLGASGSGS<br>VSSIAQGLEWAGNNGMHVANLSLGSPSPSATLEQAVNSATSRGVLVVAAS<br>GNSGAGSISYPARYANAMAVGATDQNNNRASFSQYGAGLDIVAPGVNVQ<br>STYPGSTYASLNGTSMATPHVAGAAALVKQKNPSWSNVQIRNHLKNTATS<br>LGSTNLYGSGLVNAEAATR |
| | SPU21234.1 | AQSVPWGISRVQAPAAHNRGLTGSGVKVAVLDTGISTHPDLNIRGGASFVP<br>GEPSTQDGNGHGTHVAGTIAALNNSIGVLGVAPSAELYAVKVLGADGRGA<br>ISSIAQGLEWAGNNGMHVANLSLGSPSPSATLEQAVNSATSRGVLVVAASG<br>NSGASSISYPARYANAMAVGATDQNNNRASFSQYGAGLDIVAPGVNVQST<br>YPGSTYASLNGTSMATPHVAGAAALVKQKNPSWSNVQIRNHLKNTATSLG<br>STNLYGSGLVNAEAATR |
| | WP_0532580<br>10.1 | MGALTWIRGFNGTVGRLAPRTVASKLRRTFMTPRNLPPRDWELPLLAQSE<br>RITLRFGLSALRWGQGPAVLLMHGWEGRPTQFASLITALVDNGYSVIALDG<br>PAHGRSPGREAHVLLFARAMLEAAAELPPLQAVVGHSMGGASALLAVQL<br>GLRTEALVSIAAPSRFLDVLRGFAGMVGLPARARAAFIREVELTFGMPLKH<br>LDVAHYQMNLPGLIVHAEDDTFVPVKASQAIHDAWFDSRLLRLEQGGHQK<br>VLADPRVIDGVLTLLAGCRLQARQTA |

| | | |
|---|---|---|
| | WP_099525409.1 | MNQMTWVRGVNATLGRVAPQLIASRLRERFMTPRTQPPRDWELPLLASAE RITLRFGLSALRWGSGPTVLLMHGWEGRPTQFALLIRGLVDAGYGVIALDA PAHGRSPGREANVVLFARALLEAASELPPLRAVIGHSMGGASALLATQMG LRCETLVTVAAPSRILGLLRGFARFMGLPAEARAHFVRAVETTAGIPAAHL DVQRYQLDLPGLIVHAEDDQVVPVGEADLIHKAWFDSQLLRLPAGGHQRL LSDPLLLQAVLELLEQVPQASLKALAS |
| | WP_054063832.1 | MNTLRWIRGINGTLGRVAPRVAASRMRQVFMTPRERSPRDWELPLLATAE RITLRFGLSALRWGQGPTVLLMHGWEGRPTQFASLIEALVAAGYTAVALD GPAHGQSPGHEANVVAFARALLEAAAELPPLKAVIGHSMGGASAMLATQL GLRTEALVSIAAPARVLGVLRGFAQHVGLPPRARSAFIREVERDVGMRAEH LDIGHYQMDMPGLIVHAEDDQLVAVDESRRIHEAWFDSRLLRLESGGHQR VLADPRLIDGVLALLAGRSMAQRQSA |
| | WP_071488109.1 | MGTLTWVRRFNSTLGHLVPQTVANRMRRAFMTPRELPPRDWELPLLAQSE RITLRFGLSALRWGQGPAVLLMHGWEGRPTQFASLISALVDNGYSVIALDG PAHGRSPGREAHVLLFARAMLEAAAELPPLHAVIGHSMGGASAMLAVQLG LRTEALVSIAAPSRFLDVLRGFTRMVGLPARARSAFIQEVELTFGMPLKHLD VAHYQMNIPGLIVHAEDDTFVPVKASQAIHEAWFDSRLLRLEQGGHQKVL ADPRVIDAVLALLAGCRLQERQSA |
| | WP_071486326.1 | MGALTWIRGFNGTVGRLAPHTVASKMRRTFMTPRDLPPRDWELPLLAQSE RITLRFGLSALRWGQGPAVLLMHGWEGRPTQFASLITALVDNGYSVIALDG PAHGRSPGREAHVLLFARAMLEAAAELPPLHAVVGHSMGGASAMLAVQL GLRTQALVSIAAPSRFLDVLRGFAGMVGLPPRARSAFIQEVELTFGMPLKH LDVAHYQMNIPGLIVHAEDDTFVPVKASQQIHDTWFDSRLLRLEQGGHQK VLADPRVIDAVLALLAGRRLQERQTA |
| | WP_125881520.1 | MNQMTWVRGVNATLGRVAPQLIASRLRERFMTPRTQPPRDWELPLLASAE RITLRFGLSALRWGSGPTVLLMHGWEGRPTQFALLIRGLVDAGYGVIALDA PAHGRSPGREANVVLFARALLEAASELPPLRAVIGHSMGGASALLATQMG LRCETLVTVAAPSRILGLLRGFARFMGLPAEARAHFVRAVETTAGIPAAHL DVQRYQLDLPGLIVHAEDDQVVPVGEADLIHKAWFDSQLLRLPAGGHQRL LSDPLLLQAVLELLEQVPQASLKAMAS |
| | WP_054921654.1 | MGALTWVRGFNGTVGRLAPHAVASKMRRTFMTPRDLPPRDWELPLLAQS ERITLRFGLSALRWGHGPAVLLMHGWEGRPTQFASLITALVDNGYSVIALD GPAHGRSPGREAHVLLFARAMLEAAAELPPLYAVVGHSMGGASAMLAVQ LGLRTQALVSIAAPSRFLDVLRGFTRMVGLPPRARSAFIQEVELSMGMPLK HLDVAHYQLNIPGLIVHAEDDTFVPVRAAQAIHEAWFDSRLLRLEQGGHQ KVLADPQVIDAVLALLAGCRLQERQSA |
| | WP_169851737.1 | MSTLSWIRRVNGTVGRLAPQTIANQMRRAFMTPRDLPPRDWELPLLAQAE RVTLRFGLSALRWGQGPAVLMMHGWEGRPTQFASLIDALVGAGYSVIALD GPAHGRSPGREAHVVLFARAMLEAAAELPPLHAVVGHSMGGASAMLAIQ LGLRTNALVSIAAPSRLLDVLRGFAGVVGMPARARAAFIQEVEYSLGIPLK HLDVAHYQMNIPGLIVHAEDDTFVPVKASQMIHEAWFDSRLLRLEQGGHQ KVLADPRVVEGVLALLAGCREPTRQTA |
| | WP_058426895.1 | MGTLTWIRRFNGTLGHLAPQTVANRMRRAFMTPRDLPLRDWELPLLAQSE RITLRFGLSALRWGQGPAVLLMHGWEGRPTQFASLISALVDNGYSVIALDG PAHGRSPGREAHVLLFARAMLEAAAELPPLHAVIGHSMGGASAMLAVQLG LRTEALVSIAAPSRFLDVLRGFTKMVGLPARARSAFIQEVELTFGMPLKHLD VAHYQMNIPGLIVHAEDDTFVPVKASQAIHEAWFDSRLLRLEQGGHQKVL ADPRVIDGVLALLAGARLQERQTA |
| | WP_003192569.1 | MGALTWVRGFNGTVGRLAPHAVASKMRRTFMTPRDLPPRDWELPLLAQS ERITLRFGLSALRWGHGPAVLLMHGWEGRPTQFASLITALVDNGYSVIALD GPAHGRSPGREAHVLLFARAMLEAAAELPPLYAVVGHSMGGASAMLAVQ LGLRTQALVSIAAPSRFLDVLRGFTRMVGLPPRARSAFIQEVELSMGMPLK |

| | | |
|---|---|---|
| | | HLDVAHYQMNIPGLIVHAEDDTFVPVRAAQAIHEAWFDSRLLRLEQGGHQ KVLADPQVIDAVLALLAGCRLQERQSA |
| | WP_060765121.1 | MGALTWVRGFNGTVGRLAPHAVASKMRRTFMTPRDLPPRDWELPLLAQS ERITLRFGLSALRWGHGPAVLLMHGWEGRPTQFASLIAALVDNGYSVIALD GPAHGRSPGREAHVLLFARAMLEAAAELPPLYAVVGHSMGGASAMLAVQ LGLRTQALVSIAAPSRFLDVLRGFTRMVGLPPRARSAFIQEVELSMGMPLK HLDVAHYQLNIPGLIVHAEDDTFVPVRAAQAIHEAWFDSRLLRLEQGGHQ KVLADPQVIDAVLALLAGCRLQERQSA |
| | WP_154743290.1 | MSTFKWTRGVNGALGRLAPQIIASKMRRVFMTPRNFPPRDWELPLLAQSE RITLRFGLSALRWGQGPTVLLMHGWEGRPTQFASLISALVGAGYSVIALEG PAHGRSPGREAHVLLFARAMLEAAAELPPLHAVIGHSMGGASAMLAVQLG LRTEALVSIAAPSRFLDVLRGFAGMVGLPARARSAFIQEVEHAFGMPLKYL DVAHYQMNMPGLIVHAEDDTFVSVRASQVIHEAWFDSRLMRLKQGGHQK VLADPHVIKGVLALLAGCRPAQRQTA |
